# MITE infestation of germline accommodated by genome editing in *Blepharisma*

**DOI:** 10.1101/2022.05.02.489906

**Authors:** Brandon Kwee Boon Seah, Minakshi Singh, Christiane Emmerich, Aditi Singh, Christian Woehle, Bruno Huettel, Adam Byerly, Naomi Stover, Mayumi Sugiura, Terue Harumoto, Estienne Carl Swart

## Abstract

During a sophisticated developmental process, ciliates excise numerous internally eliminated sequences (IESs) from a germline genome copy, producing a functional somatic genome. Most IESs ultimately originate from transposons but homology is obscured by sequence decay. To obtain more representative perspectives on ciliate genome editing, we assembled forty thousand IESs of *Blepharisma stoltei*, from a much earlier-diverging lineage than existing models. Short IESs (< 115 bp) were largely non-repetitive, with a pronounced ~10 bp length periodicity, whereas longer IESs (max 7 kbp) were non-periodic and contained abundant interspersed repeats. Contrary to current models, the *Blepharisma* germline genome encodes few transposases. Instead, its most abundant repeat (8000 copies) was a Miniature Inverted-repeat Transposable Element (MITE), apparently a deletion derivative of a germline-limited Pogo-family transposon. We propose MITEs as an important and eventually self-limiting IES source. Rather than defending germline genomes against mobile elements, we argue that transposase domestication actually facilitates junk DNA accumulation.

## Introduction

Ciliates are microbial eukaryotes that maintain separate germline and somatic genomes in each cell, housed in two types of nuclei. During the sexual life cycle, germline micronuclei (MICs) develop via a process of small RNA (sRNA)-assisted DNA elimination and DNA amplification into new somatic macronuclei (MACs), which are the site of most gene expression in vegetative cells. Germline-limited genome segments, called internally eliminated sequences (IESs), are excised during development from MIC to MAC. The MAC genome content is hence a subset of the germline MIC. Each of the few taxa studied so far has its own peculiarities. For example, typical *Paramecium* IESs are short, have unique sequence content, and are precisely excised, while *Tetrahymena* IESs are longer, more repetitive, and imprecisely excised (Arnaiz et al., 2012; Feng et al., 2017; Hamilton et al., 2016).

Ciliate IESs are thought to originate from cut-and-paste DNA transposons (Klobutcher and Herrick, 1997) (Figure 1B), because: (i) 5’-TA-3’ motifs at IES boundaries (*Euplotes*, *Paramecium*) resemble the terminal direct repeats of Tc1/Mariner-superfamily transposons (Klobutcher and Herrick, 1995); (ii) transposon-derived “domesticated” excisases are used to remove IESs (Baudry et al., 2009; Cheng et al., 2010; Nowacki et al., 2009); and (iii) intact transposons encoding transposases are mostly germline-limited (Arnaiz et al., 2012; Herrick et al., 1985; Jahn et al., 1993; Le Mouël et al., 2003). Recently, IESs with non-autonomous mobile elements that resemble miniature inverted-repeat transposable elements (MITEs) have been reported in *Paramecium* spp. (Sellis et al., 2021). MITEs are deletion derivatives of Tc1/Mariner transposons, generally short (<500 bp), lacking coding sequences, bounded by terminal repeats, and are common in plants and animals (Feschotte et al., 2002). However, the autonomous counterparts of most *Paramecium* putative MITEs, including the most abundant ones with thousands of copies, have not been identified.

**Figure 1.**
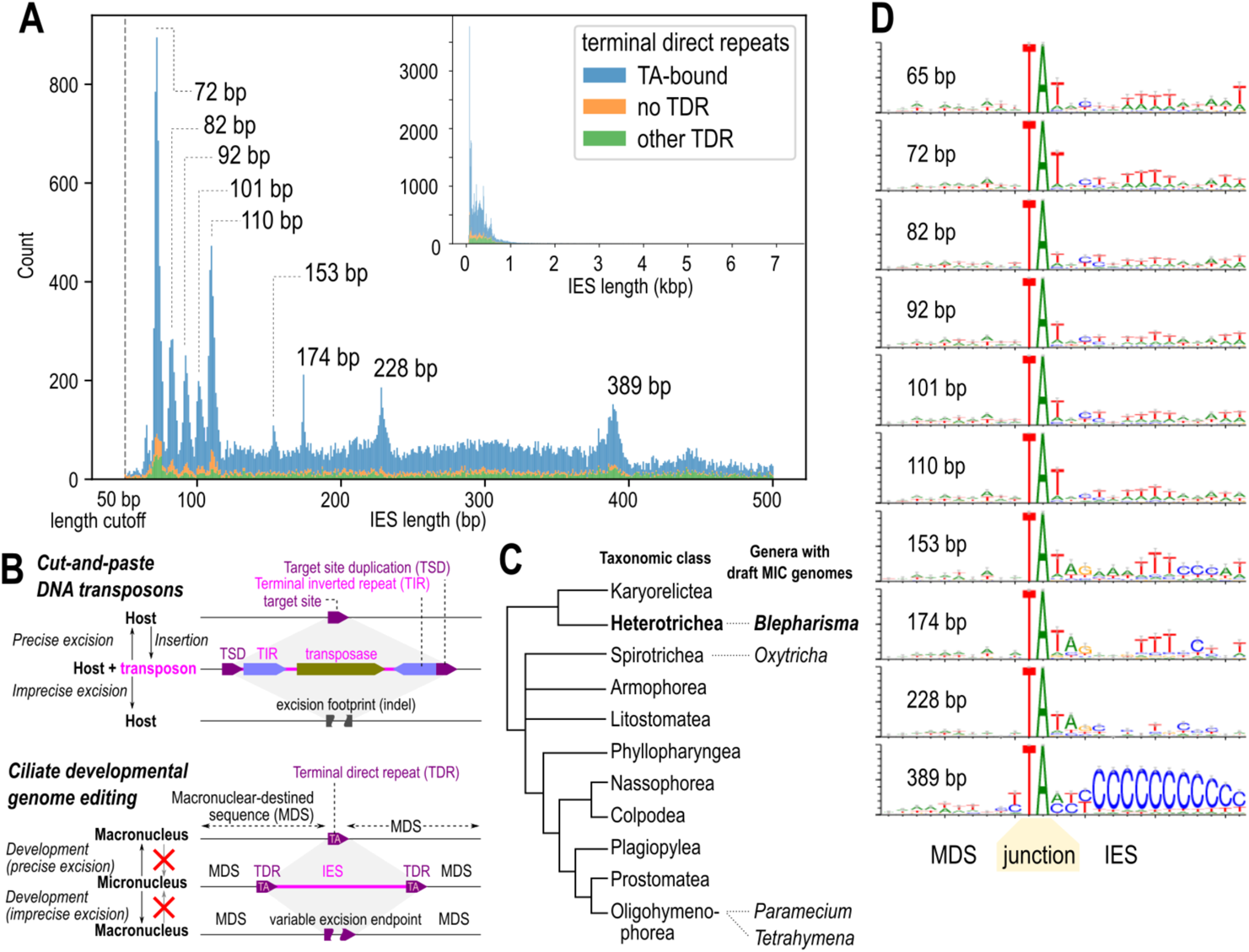
A “hybrid” IES length distribution with periodic length peaks for short IESs. (**A**) IES length histogram (0 to 500 bp (inset: full range), stacked bars for types of terminal direct repeats (TDRs) at IES boundaries. Peaks for IES size classes discussed are marked. (**B**) Comparison of cut-and-paste DNA transposons (above) and ciliate genome editing (below), showing parallels between target site duplications (TSD) of transposons and terminal direct repeats (TDRs) bounding IESs, and effects of precise vs. imprecise excision. (**C**) Diagrammatic tree of ciliates (following Lynn, 2008), branch lengths arbitrary. Genera with draft MIC genomes listed on right. (**D**) Sequence logos for MDS-IES junctions for TA-bound IESs of specific size classes, centered on the “TA”. See also Figure S1.

Developmental DNA elimination has been called “genome defense” because the process removes IESs, which not only derive from selfish genetic elements (transposons), but are often intragenic and hence deleterious if not removed (Yao et al., 2003). The “defense” analogy was popularized due to parallels to other eukaryotes where small RNA-mediated DNA heterochromatinization is thought to suppress mobile element proliferation (Coyne et al., 2012; Grewal and Jia, 2007; Vogt and Mochizuki, 2013). Ciliates have been proposed to use development-specific sRNAs to guide DNA elimination; in oligohymenophoreans, they mark sequences for elimination (Mochizuki et al., 2002; Sandoval et al., 2014; Yao et al., 2003), whereas spirotrich sRNAs mark sequences to be retained (Fang et al., 2012; Zahler et al., 2012). Histone modifications are also required for elimination (Liu et al., 2007; Taverna et al., 2002). sRNAs may not always be strictly necessary: in *Paramecium*, knockdown of key sRNA biogenesis enzymes had a smaller effect on shorter IESs, and were only weakly correlated with the more potent effects of knocking down the main IES excisase (Sandoval et al., 2014; Swart et al., 2014).

Other phenomena during genome editing vary markedly between the few model species studied in detail (reviews: (Chalker et al., 2013; Coyne et al., 2012; Rzeszutek et al., 2020)). For example, in all species, germline chromosomes are fragmented into smaller somatic ones to some degree, but spirotrichs produce extremely short somatic “nanochromosomes” with only one or a few genes. “Unscrambling” of nonsequential MAC-destined sequences into the correct order in the somatic genome occurs frequently in some spirotrichs, e.g. *Oxytricha* and *Stylonychia* (Prescott and Greslin, 1992), infrequently in *Tetrahymena* (Hamilton et al., 2016), and has not been reported in other ciliates (e.g. *Paramecium and Euplotes*). Draft-quality germline genomes are available from only two out of eleven class-level taxa (following taxonomy of Lynn, 2010): Oligohymenophorea (Arnaiz et al., 2012; Guérin et al., 2017; Hamilton et al., 2016; Sellis et al., 2021) and Spirotrichea (Chen et al., 2014) (Figure 1C).

Since it is not apparent which genome editing elements are common to all ciliates, we targeted the heterotrich *Blepharisma stoltei* (class Heterotrichea), whose last common ancestor with other ciliates with sequenced germline genomes is the last common ancestor of all ciliates (Gao and Katz, 2014). *Blepharisma* has been a laboratory model for photobiology (Giese, 1973) and mating factors (Kubota et al., 1973; Miyake and Beyer, 1974; Miyake et al., 1991; Sugiura and Harumoto, 2001), so cultivated strains and protocols for inducing conjugation and development are available, and now too an accurate, highly contiguous draft somatic genome (Singh et al., 2021). The somatic genome encodes a likely IES excisase, *Blepharisma* PiggyMac (BPgm), most closely related to the main IES excisases of *Paramecium* (PiggyMac) and *Tetrahymena* (Tpb2). Other somatic PiggyBac paralogs are also present but lack a complete “catalytic triad”, similar to the situation in *Paramecium* (Bischerour et al., 2018). BPgm is upregulated during formation of the new somatic MAC alongside other development-specific genes, including homologs of sRNA biogenesis proteins implicated in genome editing (Singh et al., 2021).

In this study, we assembled a draft germline genome for *Blepharisma stoltei*. Through single molecule long read sequencing and targeted assembly, we assembled IESs including many with long, repetitive elements, which is not feasible with short read shotgun sequencing alone. We found about ten thousand short (≤115 bp), precisely excised IESs with a periodic length distribution like *Paramecium*’s. However most IESs (about thirty thousand) were longer, up to several kbp, and, importantly, also include a Tc1/Mariner transposon whose non-autonomous MITE was also the most abundant repeat in the genome. Complementing the genomic analyses, we also identified small RNAs expressed during sexual development with characteristics of scnRNAs that guide DNA elimination in other ciliates. These results show that characteristics of germline-limited DNA in ciliates may be disjunct to phylogeny, and also illustrate how MITEs could be an intermediate stage in the origin and proliferation of IESs.

## Results

### Detection and targeted assembly of ca. forty thousand germline-limited IESs

To investigate the *Blepharisma* germline genome we enriched germline micronuclei from *B. stoltei* strain ATCC 30299, and reconstructed 39799 IESs (13.2 Mbp total, average coverage ~45x) scaffolded on the previously assembled 41 Mbp somatic genome (Singh et al., 2021) using a mapping and targeted assembly approach for PacBio long reads (Seah and Swart, 2021). This MAC-scaffolded germline assembly is here referred to as the “MAC+IES” assembly. About 70% of all predicted IESs were intragenic (within coding sequences or introns), implying precise excision of IESs, as they would otherwise cause deleterious translation frameshifts. However, genes occupied 77% of the somatic assembly (excluding telomeres), so there was a small but statistically significant (*p* = 3 × 10^-269^) relative depletion of intragenic IESs.

### A “hybrid” IES length distribution with periodic length peaks for short IESs

Most IESs were short (median 255 bp, mean 331 bp), but the distribution was long-tailed (90th percentile 603 bp, max 7251 bp). The length distribution was not unimodal, but had multiple peaks at specific length values (Figure 1A, Table S1). It appeared to be a “hybrid” distribution composed of two ranges: a “periodic” range, from ~65 to 115 bp (10778 IESs), and a “non-periodic” range, >115 bp (29021).

The “periodic” IES size range contained sharp peaks every 10 to 11 bp, similar to the periodicity of IESs in *Paramecium tetraurelia* (Arnaiz et al., 2012; Guérin et al., 2017). The first peak in *B. stoltei* was centered at 65 bp, compared to 28 bp in *P. tetraurelia*, and there was no “forbidden” peak. The most abundant “periodic” length peaks were at 72 bp and 110 bp. The “non-periodic” range (≥115 bp) contained isolated peaks at 153, 174, 228, and 389 bp, which has no obvious periodicity. Only 9701 IESs (total 1.36 Mbp) were contained within the size classes represented by the above peaks (both periodic and non-periodic) (Table S1), meaning that most IESs had lengths outside the peak values.

### IESs are bounded by heterogeneous direct and inverted terminal repeats

In other ciliates, IES boundaries often have conserved terminal repeat motifs that could reflect excisase cut site preferences or IES origins from specific classes of transposons (Klobutcher and Herrick, 1997). We therefore searched for both direct and inverted terminal repeats in *Blepharisma* IESs.

About three quarters of IESs (30212, 9.43 Mbp) were bounded by terminal direct repeats (TDRs) that contained the subsequence TA (“TA-bound”). Other non-TA TDRs accounted for another 6566 (2.85 Mbp); the remainder were not TDR-bound, though some may be assembly errors (Figure 1A). Like most ciliates, *B. stoltei* genomes were AT-rich (somatic 33.5% GC, IESs 33.3% GC) but the number of TA- and TDR-bound sequences was unlikely to be due to nucleotide composition alone (Figure 2A, 2B). The most common TDRs were simple alternations of T and A (TA, TAT/ATA, TATA), especially in IESs up to 228 bp (Figure 2C), with the exception of TAA/TTA (see below).

**Figure 2.**
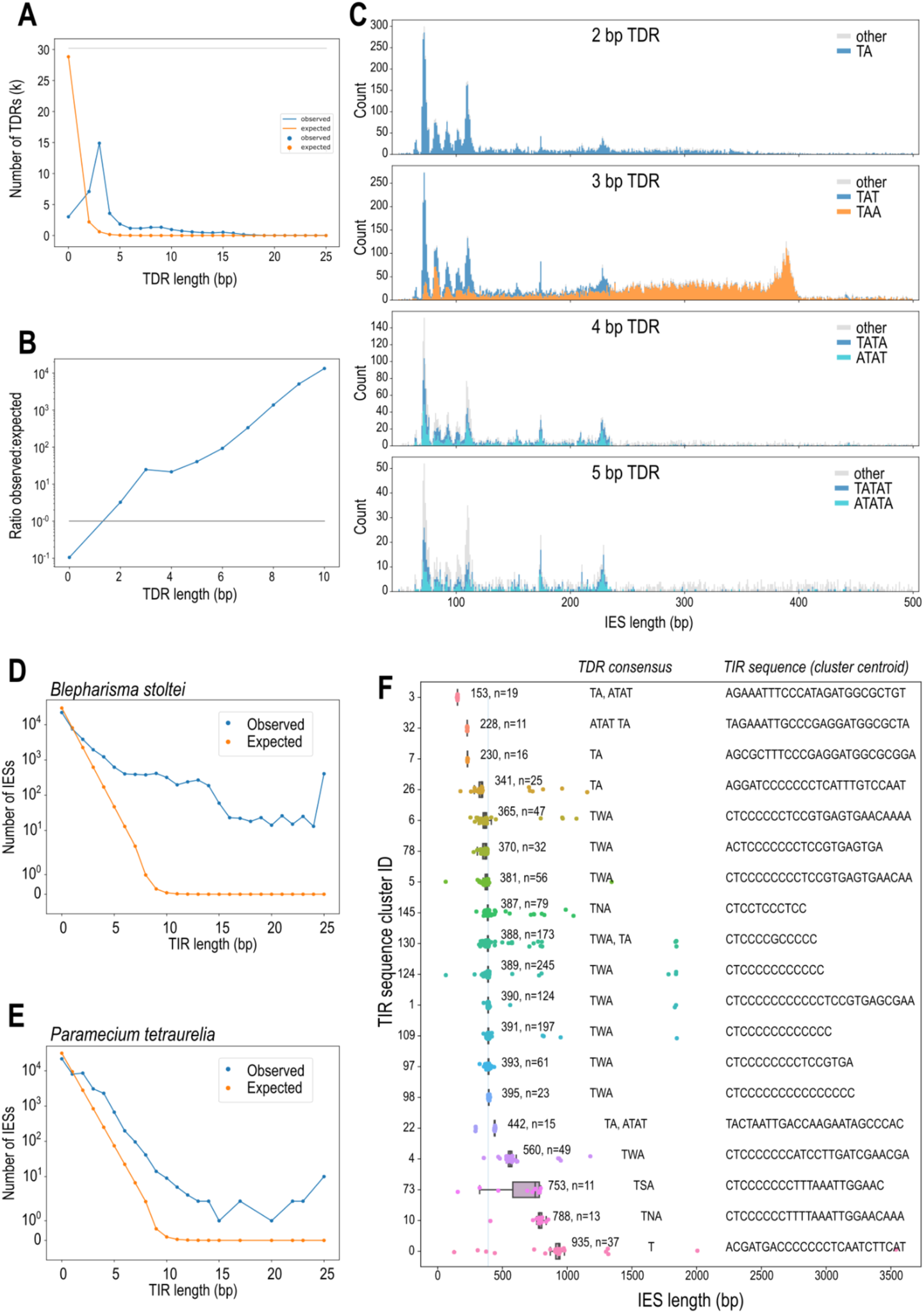
IESs are bounded by heterogeneous direct and inverted terminal repeats. (**A**) Numbers of terminal direct repeats (TDRs) per TDR length observed (blue) vs. number expected by random chance if bases were independently distributed (orange). (**B**) Ratio of observed to expected numbers of TDRs by length. (**C**) Length distributions of IESs containing TDRs of lengths 2, 3, 4, and 5 bp; the most abundant TDR sequences per TDR length are shown in color (sequences and their reverse complements are counted together, because TDRs could be encountered in either orientation, e.g. TAA/TTA), simple T/A alternations are in shades of blue. NB: plots in panel C have different vertical axis scales. (**D**) Observed IESs per terminal inverted repeat (TIR) length vs. expected number by chance alone. (**E**) Same as panel D but for *P. tetraurelia*. (**F**) Lengths (scatter-overlaid boxplot) of IESs containing long TIRs (≥10 bp), grouped by their TIR sequence (rows). Each TIR-cluster is annotated with the median IES length (bp), cluster size (n), TDR consensus sequence, and TIR representative sequence. See also Figure S2.

Erroneous, low-frequency excision of MAC-destined sequences (MDSs) by the excision machinery (“cryptic” IESs) was also detected in MAC DNA libraries, with a slight peak at 72 bp (Figure S1C). Of 10048 cryptic IESs, 56% were TA-bound; TAA/TTA-bound IESs were also common, which suggests that the observed TDRs, including TAA/TTA, represented intrinsic cut site preferences of the domesticated excisase(s) (Figure S1C to F).

Terminal inverted repeats (TIRs) at IES junctions were heterogeneous among IES size classes (Figure 1D, Figure 2F), and no single TIR motif was generally conserved across all *Blepharisma* IESs, unlike the common 5’-TAYNR-3’ motif of *Paramecium* IESs. Considering only TA-bound IESs, boundaries of “periodic” IESs had a weak consensus <u>5</u>’-T<u>A</u>T rrn ttt t-3’ (weakly conserved bases in lowercase), whereas IES from “non-periodic” peaks had other signatures, e.g. 5’<u>-</u>TAT Agn nnT TT-3’ for both ~153 and ~174 bp IESs. Despite their heterogeneity, TIRs were more common and longer than expected by chance, even with a strict criterion of no gaps or mismatches (Figure 2D to F). Sequence clustering of long (≥10 bp) TIRs showed distinct TIRs associated with specific IES lengths. Additionally, 376 completely palindromic IESs were identified, of which 153 (40.7%) fell within the same ~228 bp length peak, despite comprising several apparently unrelated palindrome sequences (Figure S2, Supplemental Information).

IESs in the ~389 bp size peak had distinctive TDRs and TIRs, suggesting they are a family of “mobile IESs”, i.e. homologous IESs inserted at nonhomologous genomic sites (Sellis et al., 2021), described further below (see “Pogo/Tigger-family transposon with abundant MITEs”).

### Repeat elements are abundant in long, non-periodic IESs

Mobile elements that have recently proliferated should appear as interspersed repeat elements in the genome. As identified by RepeatModeler, a quarter of the assembly (12.7 Mbp, 23.3%) was composed of such interspersed repeats; like in other model ciliates (Chen et al., 2014; Hamilton et al., 2016), they made up a greater proportion of germline-limited IESs (71.0%) than the somatic genome (8.12%) (Figure 3A). The majority of sequence in IESs ≥115 bp was annotated as repetitive, whereas the converse was true for shorter “periodic” IESs (Figure 3C), paralleling short IESs in *Paramecium* which are mostly unique sequences (Arnaiz et al., 2012).

**Figure 3.**
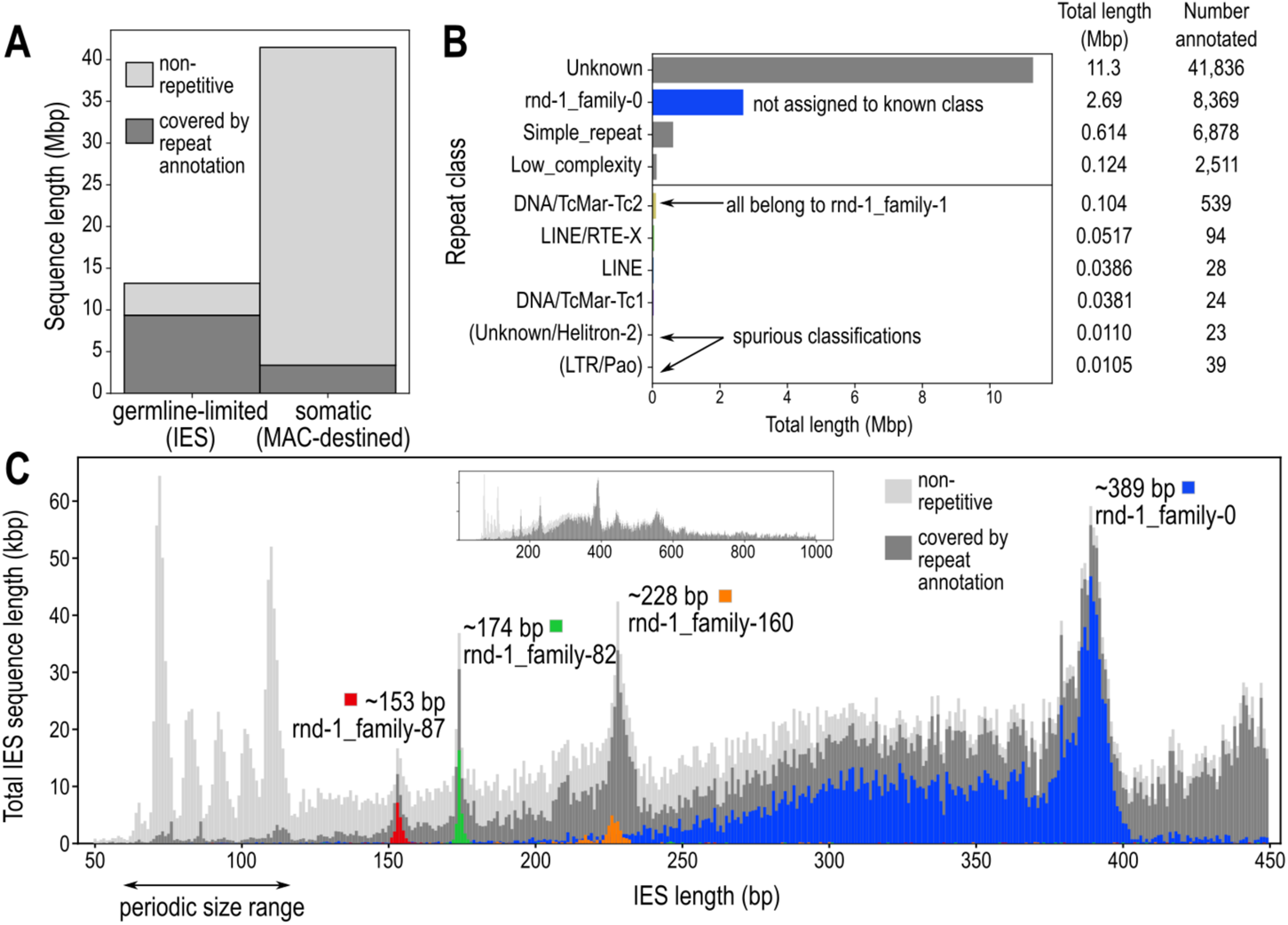
Repeat elements are abundant in long, non-periodic IESs. (**A**) Total sequence length annotated as interspersed repeats vs. non-repetitive, in germline-limited vs. somatic parts of the genome. (**B**) Classification of repeat families by RepeatClassifier, and total annotated length per repeat class. (**C**) Total sequence length (vertical axis) per IES size class (horizontal axis), stacked plot of non-repetitive fraction vs. interspersed repeats, with the most abundant repeat families in the four non-periodic peaks overlaid in color. Inset: Distribution to 1000 bp. See also Figure S3.

Most interspersed repeats could not be classified to a known transposable element class by RepeatClassifier (Figure 3B, Table S2). The most abundant classifiable type was “DNA/TcMar-Tc2”, all of which actually belonged to a single repeat family rnd-1_family-1, followed by “LINE/RTE-X”. The most abundant family, rnd-1_family-0, was unclassified and made up 21.2% (2.69 Mbp) of total repeats. Families rnd-1_family-0 and rnd-1_family-1 were related to each other and are discussed further below (“Pogo/Tigger-family transposon with abundant MITEs”).

Three non-periodic IES length peaks (153, 174, 389 bp) could be attributed to specific repeat families, suggesting that they proliferated recently (Table S3, Figure 3C, S3B). This was most pronounced for the ~389 bp peak, where 68.5% of the sequence content belonged to rnd-1_family-0, whereas about a quarter of the ~153 and ~174 bp peaks was composed of repeat families rnd-1_family-87 (palindromic) and rnd-1_family-82 respectively.

### Germline-limited repeats include few autonomous transposons but many MITEs

Unlike *Tetrahymena* and *Oxytricha* where transposases are abundant in the germline-limited IESs but rare in the somatic genome (Chen et al., 2014; Hamilton et al., 2016), *Blepharisma* encoded only a few dozen identifiable transposase domains in either the germline-limited or somatic genomes. Cut-and-paste DNA transposase domains of the DDE/D superfamily identified in *Blepharisma* included DDE_1 and DDE_3 (Tc1/Mariner family), DDE_Tnp_1_7 (PiggyBac), DDE_Tnp_IS1595 (Merlin), and MULE (Mutator) (Figure 4E, Table S4). Not all copies of DDE/D transposase domains in *Blepharisma* contained an intact catalytic triad, suggesting that some may be inactive fragments or pseudogenes. Nonetheless, domains with an intact triad were found in both germline-limited and somatic sequences. In general, the expression level of somatic transposase genes was substantially higher than germline-limited ones (Figure S4). This contrasts with the observations in *Oxytricha* of abundant germline-limited transposase gene expression (Chen et al., 2014).

**Figure 4.**
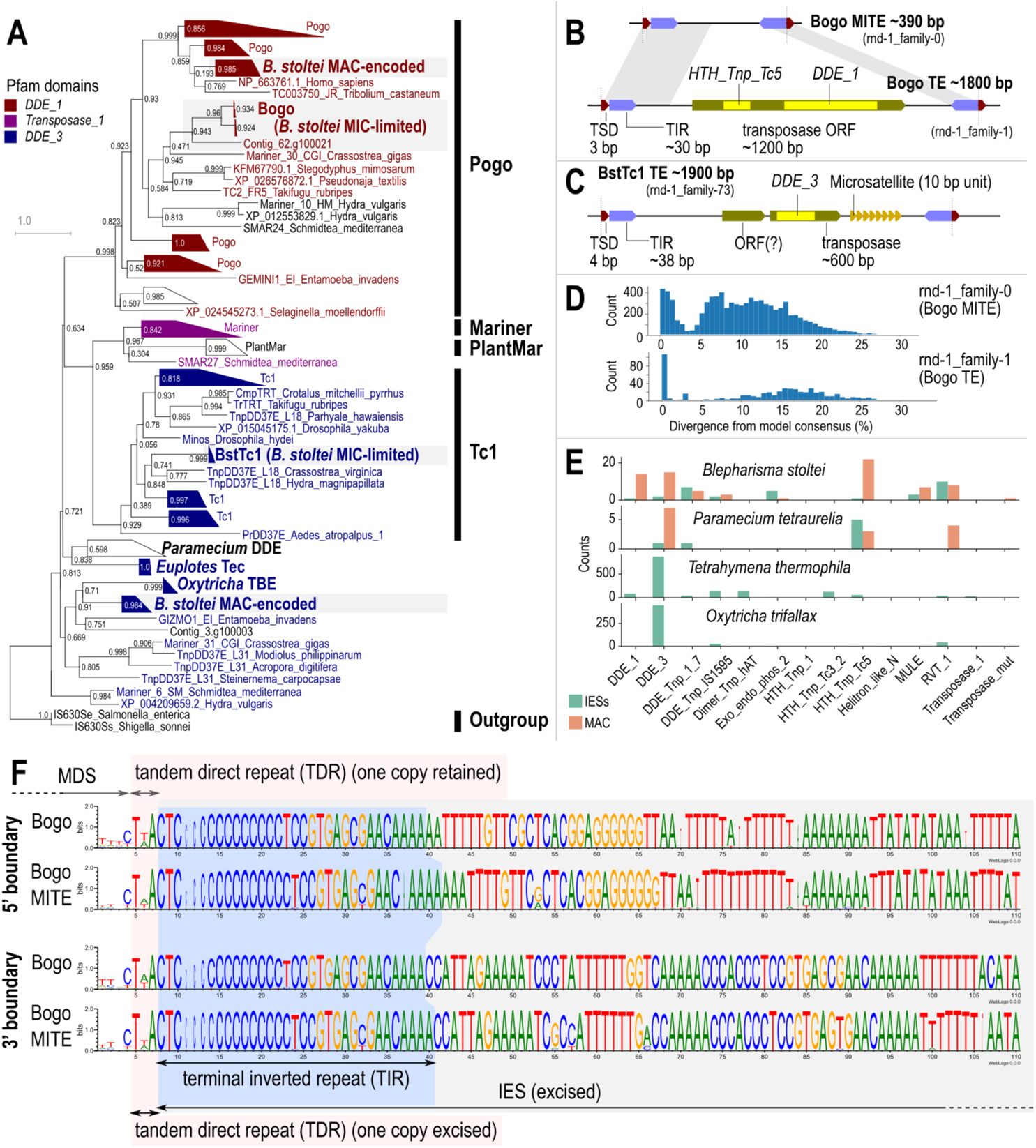
Germline-limited repeats include few autonomous transposons but many MITEs. (**A**) Phylogenetic tree of DDE/D domains for Tc1/Mariner superfamily, including *B. stoltei* germline-limited (Bogo and BstTc1) and somatic transposases. (**B**) Diagram of features in Bogo and BogoMITE; TSD – target site duplications, TIR – terminal inverted repeats, HTH_Tnp_Tc5, DDE_1 – conserved domains. (**C**) Diagram of features in BstTc1: DDE_3 – conserved domain. (**D**) Histograms of sequence divergence from repeat family consensus for copies of the Bogo and BogoMITE repeat families annotated by RepeatMasker; for rnd-1_family-1, most low-divergence copies (<5% divergence) were short fragments, but all full-length copies were low-divergence. (**E**) Counts of transposase-related domains in different ciliates from six-frame translations of somatic vs. germline-limited genome sequence. See also Figure S4. (**F**) Sequence logos for Bogo and BogoMITE repeat boundaries, aligned on the terminal inverted repeats (TIRs) and terminal direct repeats (TDRs). 3’-boundaries have been reverse complemented to show the TIRs. Sequence logos were generated from alignments of full-length, intact Bogo elements (>1.8 kbp) and BogoMITEs (between 385-395 bp), with columns comprising >90% gaps removed.

To identify intact transposon units, we examined the seven repeat families in the MAC+IES assembly classified by RepeatClassifier (Figure 3B). Of these, only two were predominantly germline-limited and represented by more than one full-length copy, namely rnd-1_family-1 and rnd-1_family-73 (Table S5). They contained distinct transposases from those found in the MAC genome (Figure 4).

#### Pogo/Tigger-family transposon with abundant MITEs

Repeat elements of rnd-1_family-1 were bound by a ~30 bp terminal inverted repeat (TIR) 5’-CTC CCC CCC CCC CTC CGT GAG CGA ACA AAA-3’ whose poly-C run length was variable, possibly from assembly errors, and were flanked by a putative target site duplication (TSD) 5’-TAA-3’ (or its reverse complement 5’-TTA-3’) (Figure 4B). All thirty intact (≥95% of consensus length) copies of this family were found within IESs, and had high sequence identity, with median 0.5% divergence from consensus.

The encoded transposase contained two domains characteristic of Pogo transposases from the Tc1/Mariner superfamily: a DDE/D superfamily endonuclease domain (Pfam domain DDE_1) and a helix-turn-helix domain (HTH_Tnp_Tc5) (Gao et al., 2020). The conserved acidic residues (“catalytic triad”) characteristic of DDE/D transposases (Yuan and Wessler, 2011) were also present, with the motif DD35D, i.e. all three residues were Asp, 35 a.a. between the second and third conserved Asp. A phylogeny of the DDE_1 domain placed the transposase in the Pogo/Tigger family, most closely related to the Tc2 subfamily and a sequence from the oyster *Crassostrea*, all of which also had the DD35D motif (Figure 4A).

The transposase appeared to be germline-limited, with only ten partial Tblastn hits in the somatic MAC genome (seven on “cruft” contigs) mostly overlapping the HTH_Tnp_Tc5 domain (17 to 84 a.a., E-values 2.3 x 10^-12^ to 1.4 x 10^-6^) and no matches to the DDE_1 domain. However, the TIR did not match previously characterized TIR signatures for the Tc2, Fot, and Pogo subfamilies. A search of all *B. stoltei* IES sequences against HMMs for known DNA transposon TIRs in the Dfam database found only three matches with E-value < 0.01, none from the above subfamilies.

The same TIR and TSD were also found in another repeat family rnd-1_family-0, which was the most abundant repeat in the genome (Figure 4F), but these were short elements without any predicted coding sequences. rnd-1_family-0 elements often constituted most of the ~389 bp IES size class (Figure 3C): the TSDs bounding the repeats (TAA/TTA) were the TDRs for most of these IESs (Figure 2C), and the C-rich TIR motif corresponded to the C-rich IES junctions (Figure 1D, Figure 2F). Copies of rnd-1_family-0 were also found nested in longer IESs, suggesting recent proliferation (Figure S3C). Degenerated or partial copies were found in shorter IESs (Figure 3C), with copies >5% divergence from consensus having median length 308 bp, vs. 388 bp for copies <5% divergence (Figure 4D).

Therefore, we interpreted rnd-1_family-1 as a new Pogo/Tigger transposon, with a non-autonomous derivative MITE, rnd-1_family-0. We propose the names Bogo for the transposon and BogoMITE for its MITE, as well as the new term “MITIES” (miniature inverted-repeat transposable internally eliminated sequences) to reflect their dual nature as MITEs and IESs. Given their palindromic nature, sequences underlying rnd-1_family-87 and rnd-1_family-160 repeats may also be MITIES.

#### Tc1-family transposon with microsatellites

Another IES-limited repeat family, rnd-1_family-73, also contained a DDE/D-type transposase coding sequence. Twenty-two copies were >80% of the consensus length with low sequence divergence (median 0.6% vs. consensus). A putative complete transposon bounded by a TSD 5’-TATA-3’ and a 38 bp TIR 5’-GTA CCC CCC CCC TCG TTT GTC GCA TTT TCT AGT TTT TT-3’ could be defined after manual curation of repeat boundaries (Figure 4C). Nine of these were mobile IESs, with the TSDs corresponding to the IES junctions. The remaining cases were nested in larger IESs alongside other repeat elements. Ten repeats also contained a microsatellite with ~5 to 42 copies of its 10 bp repeat unit 5’-GGG AAG GAC T-3’ (Figure 4C) not found elsewhere in the genome. We propose the name BstTc1 for this putative transposon.

The transposase encoded in full-length copies of BstTc1 contained a conserved DDE/D superfamily domain DDE_3, phylogenetically affiliated to the Tc1 family although the exact placement is unclear, grouping with only moderate support with Tc1 elements from *Crassostrea* and *Hydra* (Figure 4A). Its catalytic triad motif DD34E differed from previously reported motifs for the Tc1 family, DD41D, DD37D or DD36E (Dupeyron et al., 2020), so it may be a novel subfamily.

### Non-LTR retrotransposon sequences in both the somatic and germline genomes

Three retrotransposon repeat families in the MAC+IES assembly were classified by RepeatClassifier, i.e. “LINE” or “LINE/RTE-X” (Table S5). Two of these were more closely related with numerous very high identity sequences (>97%) (Figure 5A), suggesting recent radiation of two related retrotransposon elements, while the third was more divergent (Figure 5B; Supplemental Information). Unlike the Bogo and BstTC1-derived elements, more retrotransposon-derived sequences were detected in the *B. stoltei* somatic MAC genome than in assembled IESs (Figure 4E, Table S5). However genes in IESs may be undercounted because of (i) lower completeness of germline vs. somatic assembly; (ii) indels caused by the lower accuracy of the uncorrected long reads used to assemble IESs that prevent prediction; and (iii) shorter total length of IESs than somatic sequence.

**Figure 5.**
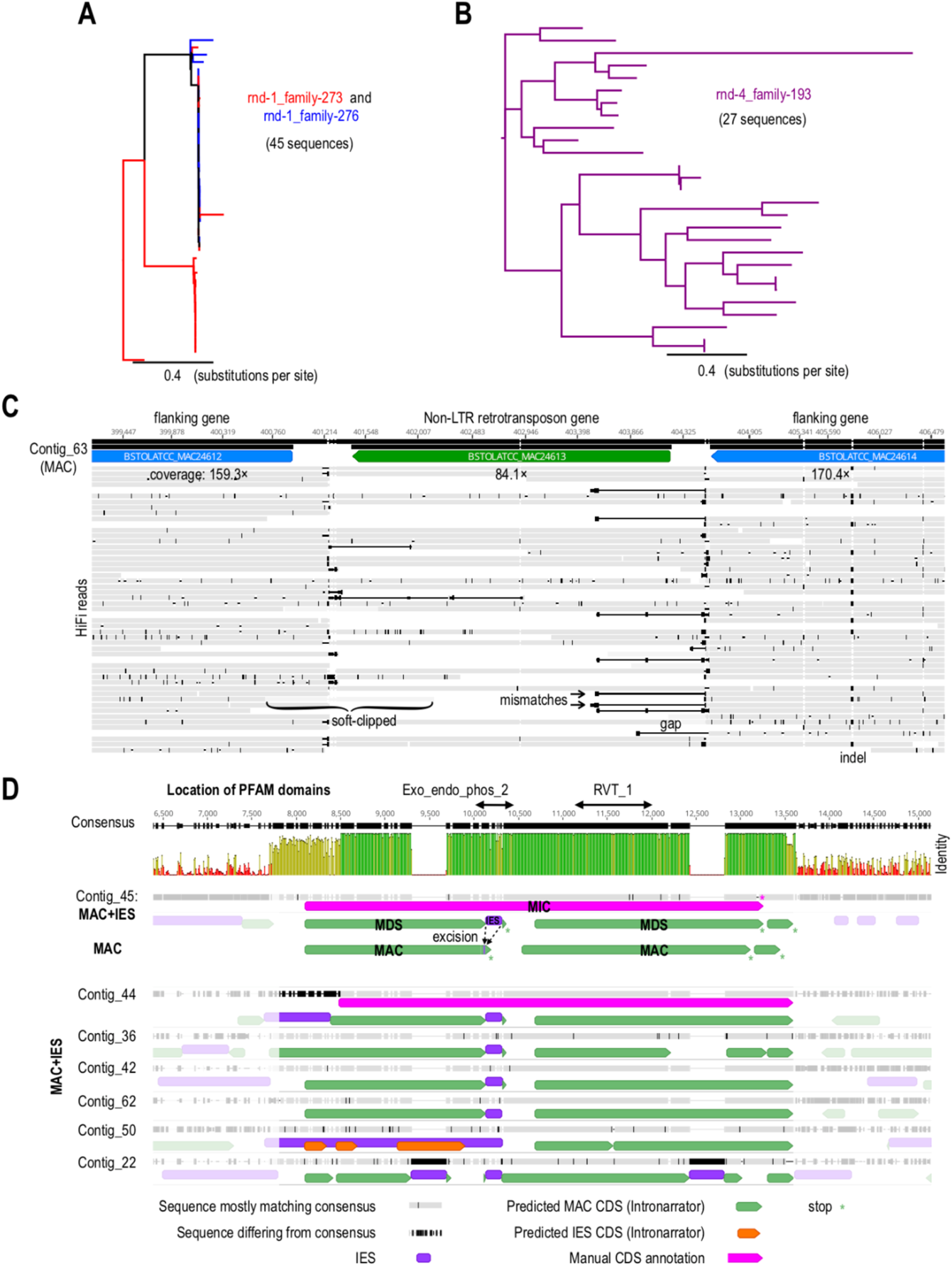
Non-LTR retrotransposon sequences in both somatic and germline genomes. (**A**) Phylogeny of rnd-1_family-273 and rnd-1_family-276 retrotransposon sequences. (**B**) Phylogeny of rnd-4_family-193 retrotransposon sequences. (**C**) Window of mapped HiFi reads from sucrose gradient-purified MACs (grey) spanning a retrotransposon gene with both an AP endonuclease domain and a reverse transcriptase domain (from rnd-4_family-193). Only sequence columns with < 90% gaps are shown. (**D**) Multiple sequence alignment of non-LTR retrotransposon copies from rnd-1_family-273. Schematic for consequences of IES excision (Contig_45). Identity scale: green=100%; gold=30-99.9%; red=0-29.9%. See also Figure S5.

Consistent with them being true somatic sequences, mappings of error-corrected long reads from a MAC-enrichment library spanned well into flanking regions (Figure 5C; Figure S5A, S5B). In each repeat family, some loci showed sharp dips in coverage suggesting partial excision as IESs while other loci did not (Figure S5B). In MAC-enriched DNA, coverage of such sequences is well above residual IES coverage (Figure S1B).

Twenty-nine genes in the main somatic assembly encoded full or partial copies of reverse transcriptase domain RVT_1 (Singh et al., 2021). The four longest retrotransposon genes also encoded an N-terminal apurinic/apyrimidinic endonuclease (Exo_endo_phos_2) domain upstream of RVT_1. This domain pair is characteristic of some proteins from non-LTR retrotransposons/LINE-like transposable elements, e.g. the BS element from *Drosophila melanogaster* (UniProt Q95SX7) (Han, 2010; Udomkit et al., 1995). In contrast to the development-specific upregulation of retrotransposon genes in *Tetrahymena* (Fillingham et al., 2004) and *Oxytricha* (Chen et al., 2014), expression of *Blepharisma* genes encoding proteins containing RVT_1 or Exo_endo_phos_2 domains was negligible in starved cells and throughout a post-conjugation developmental time series, for both germline-limited and somatic copies (Figure S4) (Singh et al., 2021). The only exception was a somatic APEX1 protein homolog (BSTOLATCC_MAC3189). APEX1 is involved in DNA repair (Fritz, 2000), and Blastp best matches of this *Blepharisma* protein to GenBank’s NR database are other similarly annotated proteins.

Six retrotransposon-derived sequences from repeat family rnd-1_family-273 contained a central IES that encoded almost half the amino acids of an Exo_endo_phos_2 endonuclease domain (Figure 5D). Excision of the IES during development thus knocks out the endonuclease domain in the somatic version of the gene. Furthermore, the repeat units as a whole had >99% identity to each other over their ~4.1 kbp length, and were flanked by dissimilar sequences (Figure 5D). The similar lengths of these IESs (173 to 182 bp), their homologous location relative to the coding sequence, and their high sequence identity (>96%) all point to a replication of an ancestral retrotransposon which coincidentally contained a sequence recognized and excised as an IES. In two of these cases, the endonuclease and reverse transcriptase domains can be linked into a single reading frame when the IES is present (Figure 5D). None of *Blepharisma*’s putative domesticated transposases are anywhere near as abundant as the retrotransposon repeats in the somatic genome, let alone show signs of substantial recent replication.

### Development-specific 24 nt small RNAs are likely scnRNAs in Blepharisma stoltei

Small RNA (sRNA) libraries were sequenced from a developmental time series, where two complementary mating types of *B. stoltei* (strains ATCC 30299 and HT-IV) were separately gamone-treated and mixed to initiate conjugation. Expression patterns of somatic genes from mRNA-seq and the morphological staging have been reported in our sister report on the MAC genome (Singh et al., 2021). Briefly: after mating types were mixed (0 h), cells paired, produced gametic nuclei by meiosis and exchanged them (2 to 18 h), followed by karyogamy (18 to 22 h) and development of the zygotic nuclei to new macronuclei (22 h onwards). At 38 h, about a third of observed cells were exconjugants.

The most abundant sRNA length classes were 22 and 24 nt, comprising 32% and 30% of the total reads respectively (Figure 6A). This is consistent with model ciliates, where Dicer-generated, mRNA-derived siRNAs employed in gene silencing are typically 21 or 22 nt long, whereas development-specific sRNAs are distinct and consistently ≥2 bp longer (Lepère et al., 2009; Mochizuki et al., 2002).

**Figure 6.**
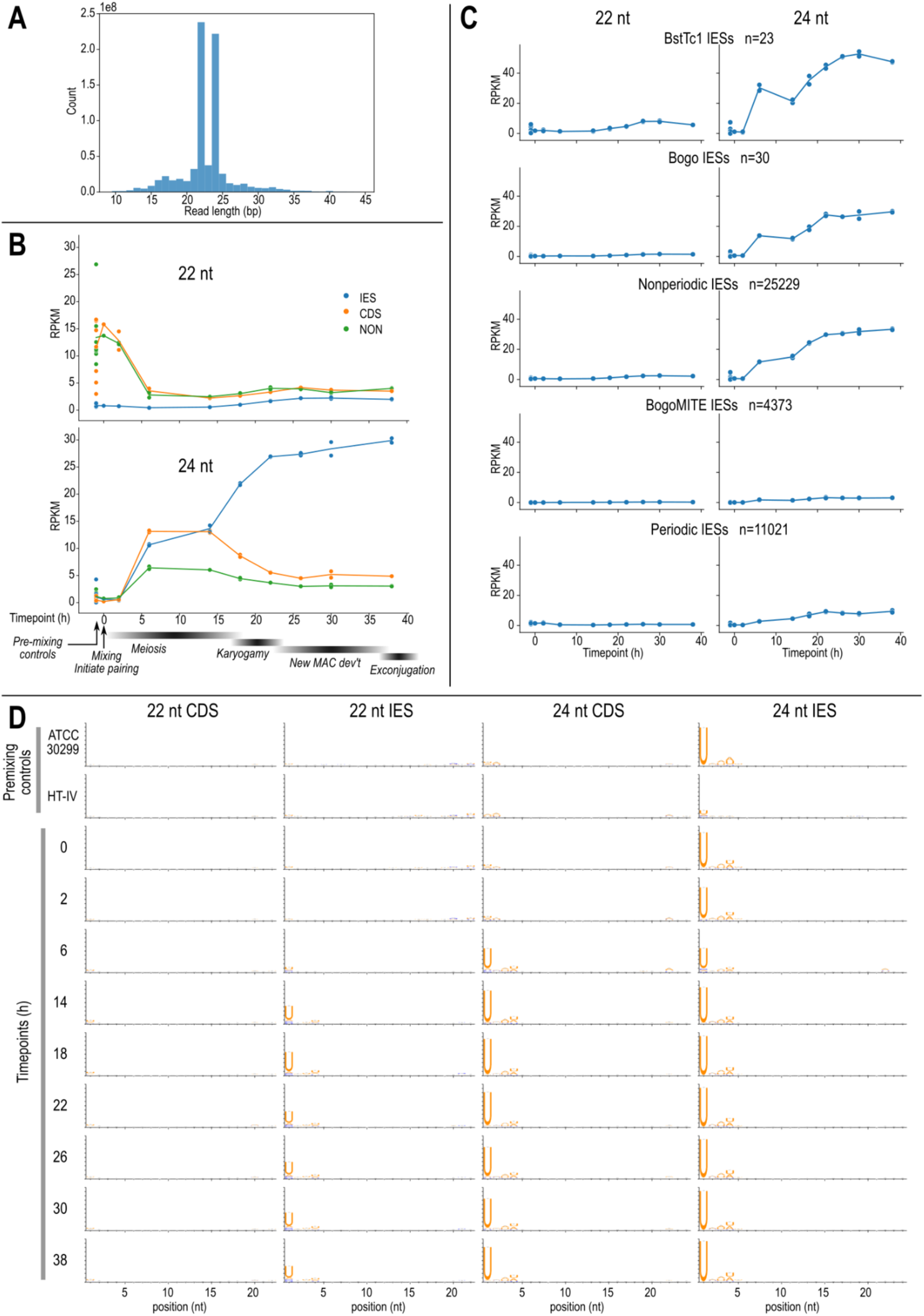
Development-specific 24 nt small RNAs are likely scnRNAs in *B. stoltei*. (**A**) Read length histogram for all sRNAs in the time series. (**B**) Relative expression (RPKM units, vertical axis) of 22 and 24 nt sRNAs mapping to different feature types across time series: blue - IES, orange - CDS, green - all other regions not annotated as IES or CDS (including UTRs and intergenic regions which are difficult to delimit exactly with available data). Timing of developmental stages inferred from morphology are labeled below (Singh et al., 2021). (**C**) Relative expression of 22 and 24 nt sRNAs mapping to different categories of IESs: containing full-length copies of BstTc1 and Bogo transposons, at least 90% covered by BogoMITE elements, IESs in the periodic length range (< 115 bp), and all other IESs (“non-periodic”). (**D**) Sequence logos for 22 and 24 sRNAs mapping to CDS and IES features in controls and different time points (rows). See also Figure S6.

Developmental dynamics of the 24 nt *Blepharisma* sRNAs resembled scnRNAs of other species. Coverage of 24 nt sRNAs mapping to all feature types initially increased from 2 to 6 h and plateaued until 14 h. Coverage over IESs increased further from 14 h to 22 h, reaching ~25 RPKM by the last time point (38 h), whereas coverage declined over coding sequences (CDSs) and other genomic regions (“NON”) after 14 h. The initial increase across all feature types coincided with meiotic stages iv to viii of (Miyake et al., 1991) (Singh et al., 2021), whereas the divergence between IESs and the rest of the genome corresponded to the onset of karyogamy (Figure 6B). In contrast, 22 nt sRNAs were initially abundant (albeit with high variance) at CDS and NON regions but low (<1 RPKM) at IESs, and declined sharply to <5 RPKM in all features from 6 h onwards (Figure 6B).

*Blepharisma* 24 nt sRNAs had a strongly conserved 5’-U base preference, like scnRNAs in other ciliates (Lepère et al., 2009; Mochizuki and Kurth, 2013; Zahler et al., 2012). For 24 nt sRNAs mapping to IESs, all time points showed conserved 5’-U except for a slight decrease at 6 h (Figure 6D, S6). 24 nt sRNAs mapping to CDSs only showed 5’-U bias after 6 h. We interpret this to mean that 24 nt sRNAs mapping to IESs were predominantly scnRNAs at all time points, whereas those mapping to CDSs initially comprised siRNAs and other types of small RNAs, before being dominated by scnRNAs from 6 h onwards. In contrast, 22 nt sRNAs mapping to CDSs showed no base biases at any time point, whereas 22 nt reads mapping to IESs had a moderate 5’-U bias only from 6 h onwards. The latter may represent true 22 nt scnRNAs, or fragments of originally 24 nt scnRNAs.

### Putative scnRNAs have lower coverage over periodic IESs and BogoMITE IESs

Relative expression levels of putative scnRNAs differed between IES size classes. Based on the IES length distribution and repeat content, we divided IESs into five groups: (1) short “periodic” IESs (≤115 bp), (2) BogoMITEs, because that was the most abundant family, (3) IESs with full-length Bogo transposons, (4) IESs with full-length BstTc1 transposons, and (5) all other IESs (“non-periodic”). BogoMITEs and periodic IESs had lower scnRNA coverage (max ~5 and 10 RPKM respectively) compared with nonperiodic IESs (~30 RPKM). The former were comparable to or even lower than expression levels over non-IES features (Figure 6C). Nonetheless, scnRNA coverage of BogoMITEs and periodic IESs showed an initial increase then plateau, without the subsequent decline seen in non-IES regions. Bogo-containing IESs had similar scnRNA coverage to other non-periodic IESs, but BstTc1-containing IESs had higher coverage (Figure 6C).

Because of the repetitive sequence content in IESs and the short sRNA length, it is possible that the expression levels calculated could be affected by mis-mapping. We reason that such mismapping would not influence the results described above, because “periodic” IESs (group 1) had low repetitive content, whereas the transposon-containing IESs (groups 2, 3, 4) each represented a single repeat family so any mismappings would be contained within the same group and count towards the same RPKM value.

## Discussion

Despite belonging to the earliest diverging lineage of ciliates sequenced to date, the germline genome of *Blepharisma stoltei* has similarities to established model species, especially the periodic lengths of short IESs like in *Paramecium*. It also provides fresh observations, notably recent proliferation of non-autonomous MITEs that have autonomous counterparts in the same genome, and retroelements in the somatic genome. Parallels between *Paramecium* and *Blepharisma* suggest that ciliate germline characteristics may be relatively plastic over evolutionary time and not strongly phylogenetically constrained.

### Comparison to IESs in other ciliates

Most *Blepharisma* IESs are short, TA-bound, and intragenic, more similar to *Paramecium* than *Tetrahymena* or spirotrichs. The most striking parallel is the sharply periodic length distribution of short IESs with peaks every ~10 bp, coinciding with the DNA helical turn, implying that the *Blepharisma* excisase complex has similar geometric constraints as proposed for *Paramecium* (Arnaiz et al., 2012). *Blepharisma* “periodic” IESs are longer on average and do not have a “forbidden” second peak, but the last peak (~110 bp; Figure 1A) is still below the upper limit where such periodicity would be expected given the properties of DNA (Figure 7 of (Arnaiz et al., 2012)). In contrast, *Tetrahymena thermophila* has a continuous distribution (average length ~3 kbp) (Hamilton et al., 2016; Seah and Swart, 2021), while *Oxytricha trifallax* non-scrambled IESs (length ~20-100 bp) have weak periodicity (Chen et al., 2014). Periodicity is consistent with a single primary IES excisase, rather than multiple excisase families, which would smooth the length distribution.

**Figure 7.**
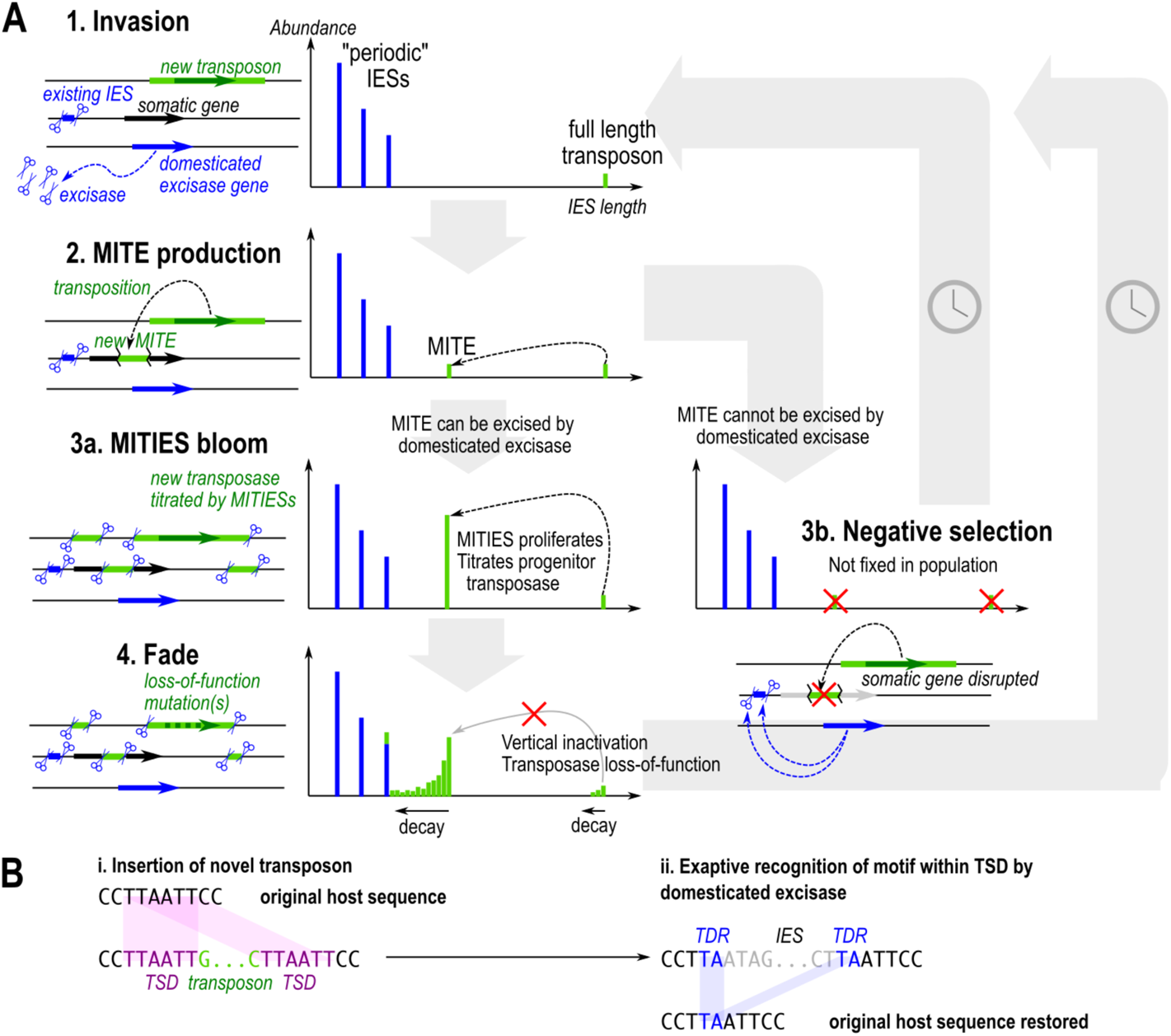
Model for IES evolution in a ciliate genome with an existing domesticated excisase. (**A**) Graphs depict IES length distribution. (**1**) Invasion of germline genome by full length transposon (green); existing IESs (blue) are excised by domesticated excisase. (**2**) New transposon produces MITIES which are both MITES and IESs. (**3a**) If MITIES can be excised by domesticated excisase, they proliferate and titrate the progenitor transposase. (**4**) Proliferation of MITIES favors vertical inactivation of the full length transposon; loss of function stops production of new MITIES, leading to eventual decay. (**3b**) If the MITE cannot be excised by domesticated excisase (i.e. it is not an IES), it is more likely to cause deleterious mutations upon insertion, and is therefore selected against and does not reach fixation. (**B**) If a transposon TSD contains a submotif that can be recognized by the domesticated excisase, it can theoretically be excised cleanly without leaving a “footprint”, avoiding potential frameshift mutations.

Longer, nonperiodic IESs of *Blepharisma* contain more repeats, including whole transposons, than short IESs. Unlike *Tetrahymena*, where 41.7% of high-confidence IESs comprise putative autonomous transposons (Hamilton et al., 2016), some of which can be grouped into families (Fillingham et al., 2004; Wuitschick et al., 2002), only a small fraction of *Blepharisma*’s long IESs encode transposases, and their length distribution is not unimodal, but long-tailed, with distinct peaks representing individual abundant families (Figure 3).

Germline-specific repeats and transposons across *Paramecium* spp. have recently been surveyed (Sellis et al., 2021), but were likely underestimated because such repeats are difficult to assemble from short-read data even with high coverage, as we saw with *Blepharisma* BogoMITE elements, (Supplemental Information, Figure S1A).

The dynamics of *Blepharisma* 24 nt sRNAs are consistent with the scnRNA turnover model, where RNA intermediates are produced from both IESs and MDSs (Malone et al., 2005; Mochizuki and Gorovsky, 2005), but those from MDSs are selectively degraded, allowing the remaining scnRNAs to mark IESs for excision. *Blepharisma* 24 nt sRNAs mapping to IESs increase more than those mapping to CDSs during post-conjugation development (Figure 6B), complementing our finding that homologs of scnRNA biogenesis proteins, Dicer-like (Dcl) and Piwi proteins, are highly upregulated during development (Singh et al., 2021).

Furthermore, higher coverage of *Blepharisma* scnRNAs in longer (presumably younger) IESs than in short (~older) periodic IESs mirrors that of *Paramecium*, where younger IESs are more likely to require scnRNAs for efficient excision (Lhuillier-Akakpo et al., 2014; Sellis et al., 2021).

The longer an IES, the more likely it will contain a promoter by chance or contain one from a transposase gene, thus giving rise to such sRNAs. This would explain the low 24 nt sRNA level of BogoMITE IESs in contrast to their autonomous counterparts (Figure 6C), though removal of both is essential. In contrast to the abundant Bogo transposon 24 nt sRNAs, expression of these and other transposase genes in RNA-seq is negligible (Figure S4). This raises the possibility that active, transcribed *Blepharisma* transposons are in fact silenced, turning most of their transcripts into 24 nt sRNAs. This is contrary to the role of scnRNAs proposed to target DNA for excision, but congruent with the role of sRNAs in transposon silencing in other eukaryotes, from which the scnRNA biosynthesis enzymes originated (Sandoval et al., 2014).

### Are MITEs a missing link in the IBAF model?

The prevailing Invasion-Bloom-Abdication-Fade (IBAF) model for the evolution of IESs hypothesizes that they originate from cut-and-paste DNA transposons that invade and proliferate (“bloom”) in the germline genome (Klobutcher and Herrick, 1997). Transposon proliferation stops (“abdication”) when its transposase is domesticated by a host promoter, releasing the transposons from purifying selection, whereupon their sequences erode by drift (“fade”). Depictions of the IBAF model usually show all the transposons expressing transposases during “bloom”, i.e. functioning as autonomous transposons (Feng and Landweber, 2021; Klobutcher and Herrick, 1997). This is reasonable for *Tetrahymena* and *Oxytricha*, which have hundreds of germline-encoded transposases that vastly outnumber those in the somatic genome (Table S4). However, *Blepharisma* and *Paramecium* only have a few dozen transposases, although germline-limited transposases may be underestimated, especially for short-read assemblies.

This discrepancy can be resolved by taking MITIESs (MITE IESs) into account. In *Blepharisma* this is best exemplified by the few autonomous Bogo transposon copies compared to thousands of non-autonomous BogoMITEs. The narrow length distribution of BogoMITEs, their high sequence identity, and occasional nested insertion inside unrelated IESs are the clearest illustrations to date of recent MITE proliferation. Bogo is also the first Pogo/Tigger transposon found in a ciliate germline genome; this subfamily is known to be especially prone to MITE formation (Feschotte and Mouchès, 2000; Guermonprez et al., 2008). The prevalence of IESs bound by terminal inverted repeats, including numerous palindromic IESs (Figure 2D, S2), also suggest many more *Blepharisma* IESs are MITE derivatives.

In *Paramecium* spp., MITEs of the Thon and Merou transposons have been identified but only numbered about a dozen copies per genome, and their transposases belong to a different transposase family than Bogo (Figure 4). The most abundant mobile IES family in *Paramecium*, FAM_2183, is probably a MITE but its autonomous counterpart was not reported (Sellis et al., 2021). MITEs as transposon/IES life cycle intermediates can hence explain why *Blepharisma* and *Paramecium* have few MIC-encoded transposases compared to *Oxytricha* and *Tetrahymena*, but nevertheless tens of thousands of IESs.

MITEs also provide a mechanism for transposon/IES proliferation self limitation (Figure 7A). When MITEs outnumber the autonomous transposon, active transposase protein is more likely to bind to target sites in MITEs than the full length transposon (“titration”), hindering the replication of the autonomous version, giving time for loss-of-function mutations to inactivate the transposases (“fade”). This “vertical inactivation” scenario (Hartl et al., 1997) was already discussed in the original IBAF proposal (Klobutcher and Herrick, 1997), but no plausible examples from ciliates were then known.

### Is “genome defense” a flawed analogy?

The IBAF model also does not explain how ciliates can consistently and precisely excise novel mobile elements from different transposon families that invade the germline genome. The domesticated excisases of *Paramecium* (Baudry et al., 2009), *Tetrahymena* (Cheng et al., 2010), and *Blepharisma* (Singh et al., 2021) belong to the PiggyBac family. Except for *Tetrahymena* Tpb2, PiggyBacs are known to perform seamless excision, where the host sequence after transposon excision is identical to that before insertion (Chen et al., 2020). This would make them the ideal progenitor for IESs within coding sequences; indeed, PiggyBac transposons are also known to produce MITEs (Mitra et al., 2013; Wang et al., 2010). By extension, the first IESs probably originated from PiggyBac transposons. But what about subsequent invasions by other transposons that leave behind “scars” upon excision? Such imprecision would cause deleterious frameshift mutations in coding regions. How can they invade the germline genome and yet avoid deleterious effects?

Part of the answer lies in the “hijacking” model proposed from *Paramecium* (Arnaiz et al., 2012; Sellis et al., 2021), whereby the domestication of PiggyBac transposase changed the dynamic for subsequent transposon invasions. New transposons would persist as IESs only if they also encode a seamless excisase, or if they can also be recognized and cut by the exapted PiggyBac transposase. The latter favors the invasion of transposons that produce a TSD containing a submotif recognized as a cut site by PiggyBac (Figure 7B). The similarity between IES and transposon boundaries would hence not be due to common origin or sequence evolution after IES fixation in the germline (Klobutcher and Herrick, 1997), but rather because of selection for transposons whose TSDs already match the excision site preferences of domesticated PiggyBac. Analogous exaptation of TSDs for excision has been demonstrated in another context: independent origin of introns from MITEs in at least two different eukaryotes, where one of the TSDs produced upon MITE insertion was co-opted as an intron splice site (Huff et al., 2016). Cross-talk between different (albeit related) transposases for MITE transposition has also been documented (Feschotte et al., 2005).

We further argue that “genome defense” is a teleological expression that confuses cause and effect. Domesticated excisases actually facilitate mobile element accumulation in the germline, by shielding them from selection by effective exclusion from the somatic genome. *Tetrahymena* is the exception that proves the rule: its domesticated excisase appears to be imprecise; correspondingly, most of its IESs are intergenic, because intragenic IESs have been efficiently removed by selection (Cheng et al., 2016; Feng et al., 2017). The origins of gene silencing by DNA methylation in vertebrates have also been reinterpreted with similar reasoning. Vertebrates have high levels of CpG methylation that inactivates transposons, which was thus proposed to “compensate for” transposon proliferation in eukaryotic genomes (Bestor, 1990). When seen from a non-teleological perspective, it is precisely because CpG-mediated transposon inactivation is so effective, preventing exposure to selection, that transposons persist, leading to larger genomes (Zhou et al., 2020).

### Why does the Blepharisma somatic genome contain retrotransposon sequences?

Transposon-related sequences are typically germline-limited in other model ciliates, which was formerly interpreted as successful “genome defense” keeping them out of the somatic MAC genome (Chen et al., 2014; Fillingham et al., 2004; Guérin et al., 2017; Hamilton et al., 2016; Swart et al., 2013). Counter to this, we found several retrotransposon-derived sequences in the *Blepharisma* MAC genome (Figure 5; Table S2). Some show signs of partial excision or possible absence of the locus in part of the population, but plenty have uniform coverage typical of somatic sequences.

Recent retrotransposon proliferation in the soma and patchy distribution of different somatic transposase classes across ciliates (Table S4) (Singh et al., 2021) suggest that “genome defense” is at best leaky. We conjecture that if foreign DNA lacks suitable target sites recognized by the excisase, it might still be marked by scnRNAs but fail to be excised or only be partially excised (e.g. the IESs in Figure 5C). Such DNA would still be deleterious if inserted intragenically.

Somatic MACs may be unable to repress mobile elements by heterochromatinization like germline MICs and other eukaryotic nuclei. In *Tetrahymena*, most MAC DNA is not associated with classical heterochromatin marks (Liu et al., 2007), while in *Paramecium* MACs, H3K27me3 is not associated with transcription repression, despite being a classic heterochromatin mark in multicellular eukaryotes (Drews et al., 2021). In such a permissive expression environment, selection against mobile elements that are not already excised as IESs may be especially effective, unless they are relatively transcriptionally inactive like the *Blepharisma* retroelements. On the other hand, regular *Blepharisma* stock culture passaging maintains a small effective population size, which would counteract selection against mobile element accumulation in the soma.

The genome defense model may lead one to dismiss IES retention in the somatic genome as excisase inefficiency or MIC contamination of the library, however, IES excision is not all-or-nothing but a continuum. Experimental evolution experiments in *Paramecium* suggest IES retention variability is itself a plastic and evolvable trait with consequences for somatic genotypic diversity (Catania et al., 2021; Vitali et al., 2019). Assembly algorithms tend to present an oversimplified, “pristine” view of somatic genomes, because they collapse repetitive and lower-coverage regions, which are characteristic of mobile elements and partially retained IESs. Accurate long read sequencing, haplotype-aware assemblers, and sequence graphs will all play a role in building a more realistic picture of somatic genome heterogeneity.

## Conclusion

Why do we credit developmental DNA elimination with defending the genome, when natural selection has been doing the hard work? Apart from technical biases during genome assembly, there is also sampling bias by using lab strains. These are often clonal and largely homozygous; if so, we would not observe accumulation of strongly deleterious foreign DNA that actually needs defending against, but only IESs that have reached fixation and that are already efficiently excised and non-deleterious. Purifying selection against deleterious IESs has had to be indirectly observed, e.g. in the lack of intragenic IESs in *Tetrahymena*, where excision is imprecise (Hamilton et al., 2016), and the statistical depletion of IES-like sequences in the *Paramecium* somatic genome (Swart et al., 2014). Similar evolutionary logic applies to prokaryotic CRISPR defense systems, where hidden fitness costs (autoimmunity) have been underestimated because those individuals are removed by selection (Stern et al., 2010), hence the phenomenon is easily misinterpreted as inheritance of acquired traits (Weiss, 2015). Most studies on ciliate developmental DNA elimination to date have focussed on the underlying molecular mechanisms, but to understand its origins and evolution we should expand our view to diverse ciliates and their germline genomes from natural populations.

## Supplemental Figure Legends

**Figure S1.**
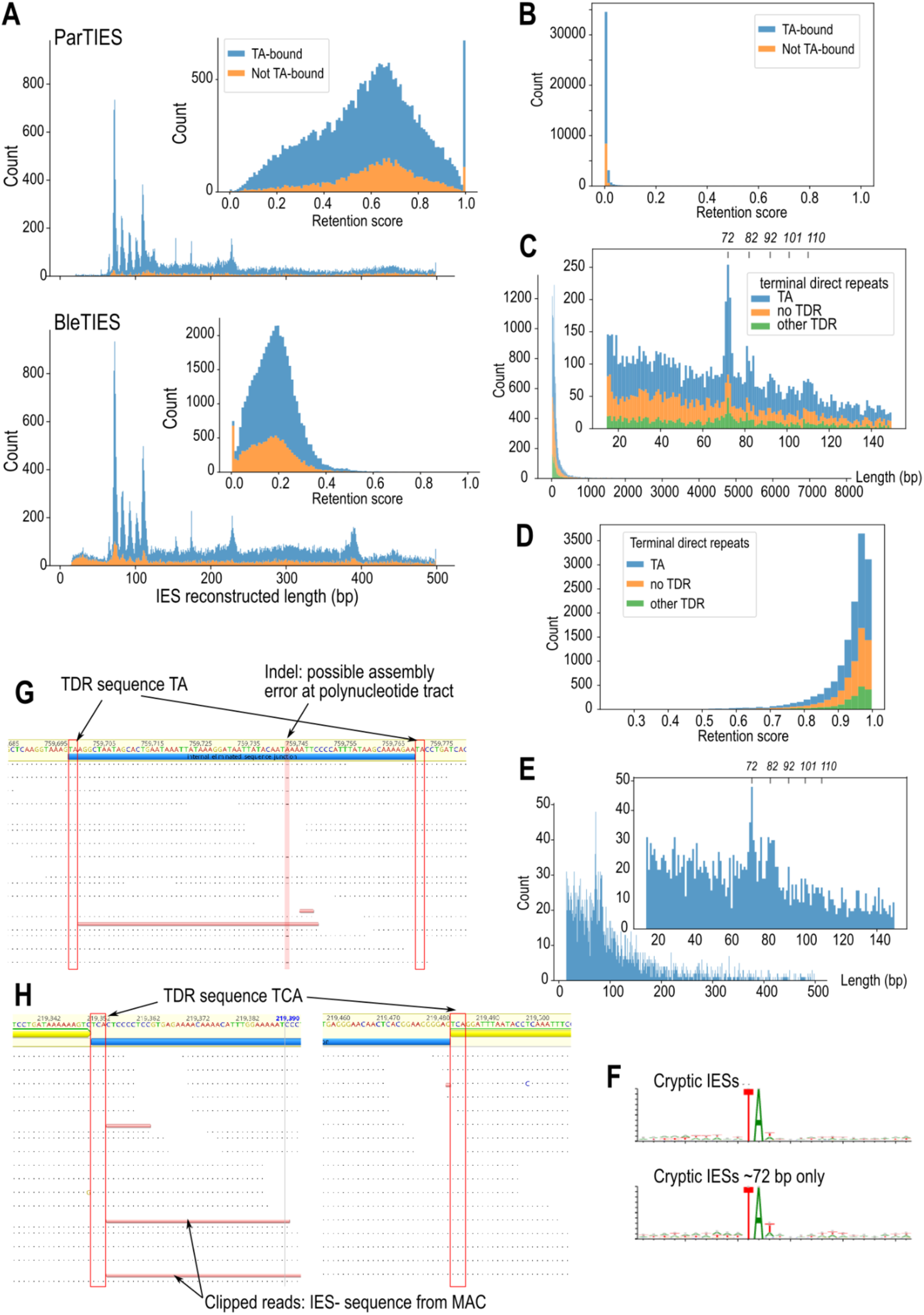
Length distributions and retention scores for different IES assembly methods, MAC library, and cryptic IESs. (**A**) Comparison of IES reconstructions from MIC-enrichment library sequenced with short reads by ParTIES (above) vs. from long reads by BleTIES (below). Main panels: IES length histograms up to 500 bp, insets: IES retention scores colored by TDR sequence type. Length peak at ~390 bp representing BogoMITE element is present in BleTIES reconstruction but not ParTIES. (**B**) Conventional IESs: retention scores computed from MAC-enriched library, sequenced with PacBio HiFi reads. (**C**) “Cryptic” IESs from MAC read library: length histogram, colored by TDR sequence type. (**D**) Retention scores of “cryptic IESs”. (**E**) Length distribution of “cryptic” IESs that contain “TTA” or “TAA” in their TDR, detail <500 bp, inset detail <150 bp. (**F**) Sequence logos of TA-bound “cryptic” IES junctions centered on the TA motif, for all cryptic IESs (above) and the subset in the ~72 bp size class (below). (**G**) Mapping pileup at IES with TA-containing TDR. For aligned reads in panels E and F, dots: bases identical to reference, dashes: gaps relative to reference, red bar: read clipping. (**H**) Mapping pileup at IES with non-TA-containing TDR.

**Figure S2.**
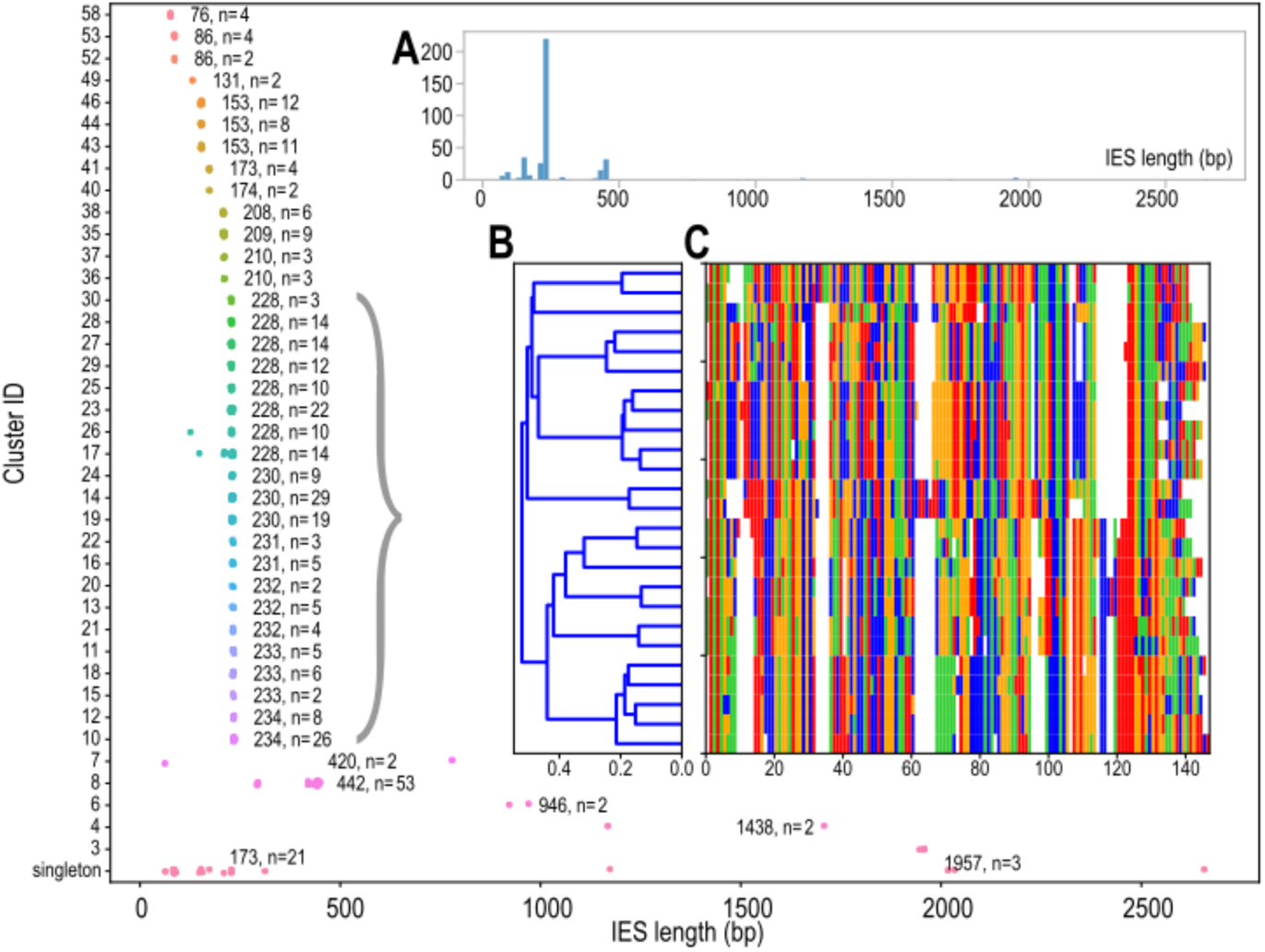
Palindromic IESs clustering and length distribution. Strip plots of IES lengths for palindromic IESs (≥90% self-alignment identity), after they have been clustered by sequence identity (rows represent clusters). Each cluster is annotated with the median IES length and the cluster size. Insets: (**A**) Overall sequence length distribution histogram for all palindromic IESs. The most common length of palindromic IESs is ~230 bp. (**B, C**) Dendrogram of sequence distance and multiple sequence alignment of palindromic IESs with ~230 bp length to illustrate that they comprise several distinct clusters of sequences.

**Figure S3.**
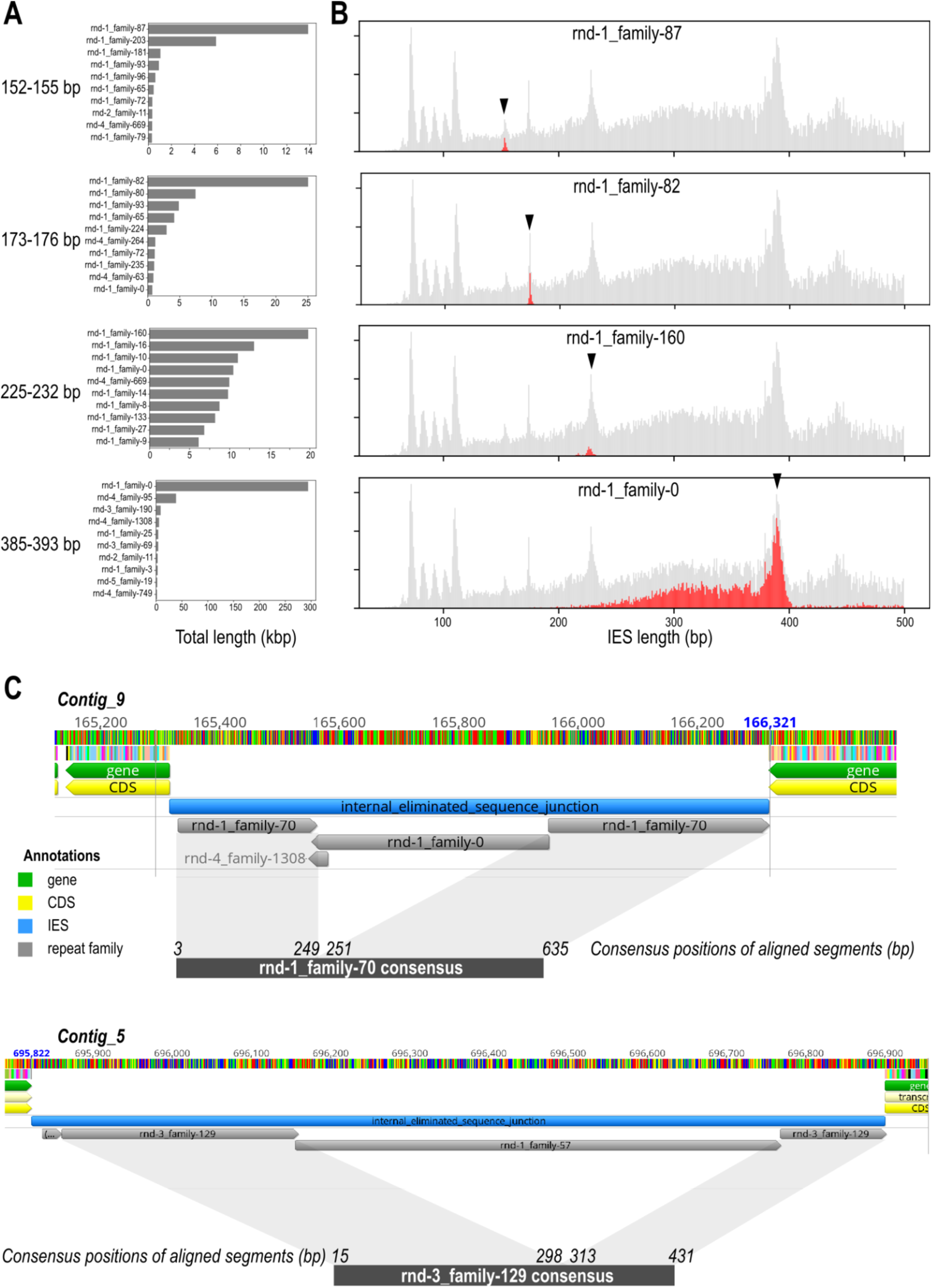
Most abundant repeat families in non-periodic IES size classes. (**A**) Total lengths (horizontal axis) of the top ten repeat families per IES size class (panel rows). (**B)** Top repeat family (by sequence length) for each IES size class (panel rows); the total length covered by that repeat family within IESs vs. the lengths of those IESs is shown in red, superimposed on the total sequence vs. IES length distribution of IESs in general (grey). Arrowheads mark centers of the size classes. (**C**) Examples of nested repeats within IESs. Nested elements can be recognized when the two outer repeat elements belong to the same family and align to consecutive parts of its family’s consensus sequence, implying that the inner element has likely been inserted into the middle of an existing element. Coordinates of the split segments are relative to the repeat family consensus.

**Figure S4.**
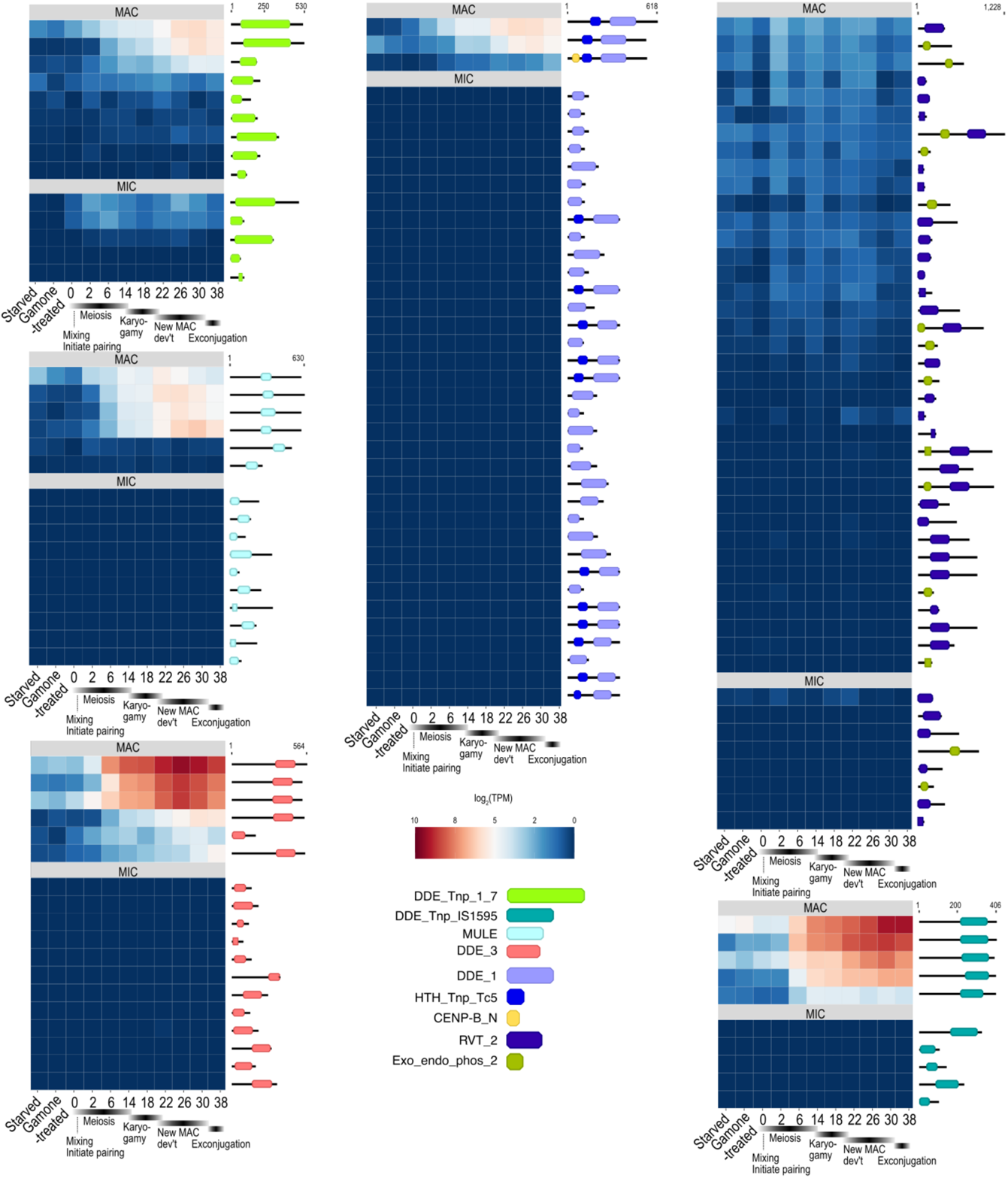
Expression of genes with transposase domains. Comparison of expression levels for MAC- vs. MIC-limited transposase-related domains across developmental time series; heatmap color scaled to log(transcripts per million). Domain architecture shown diagrammatically.

**Figure S5.**
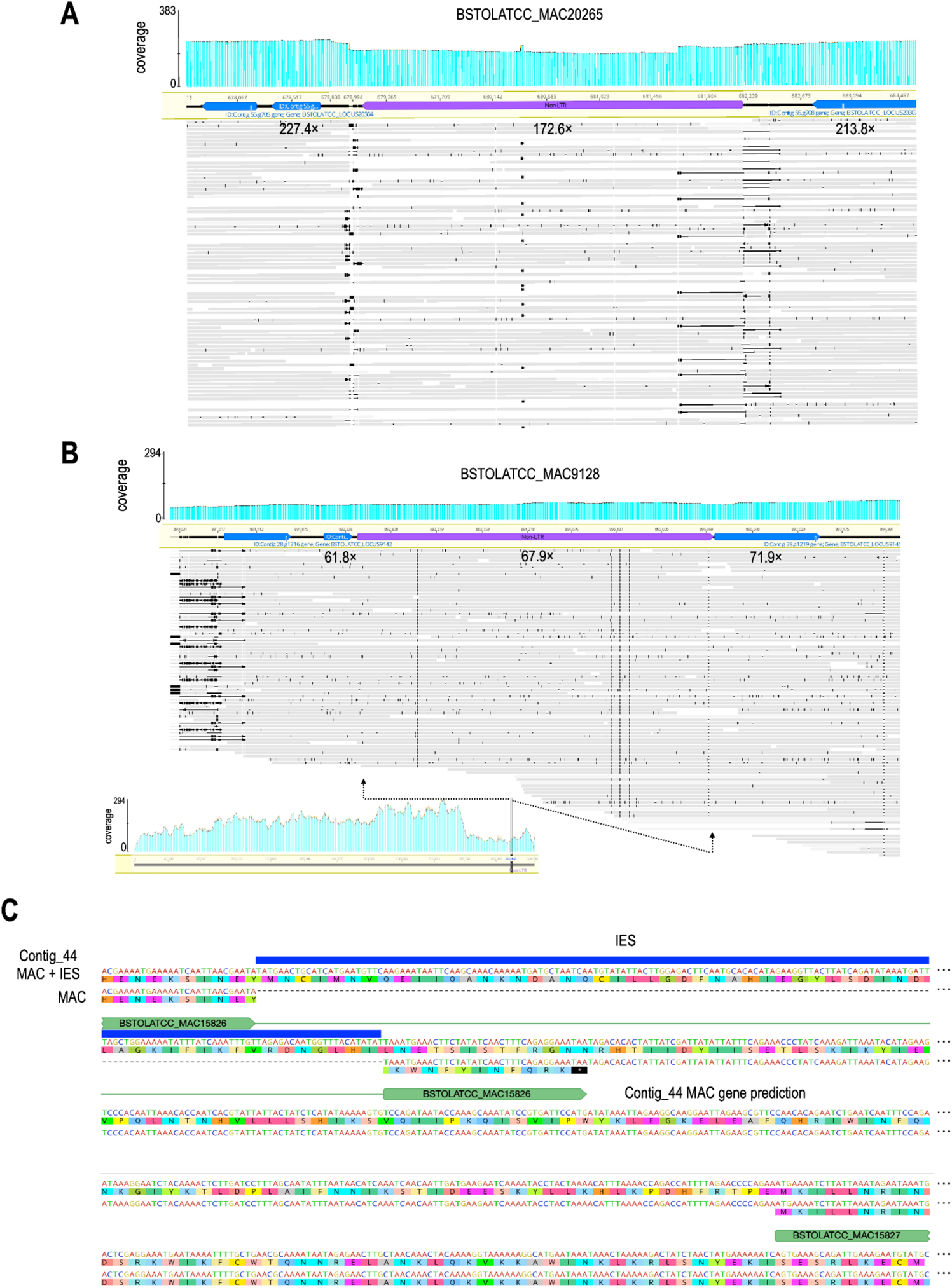
Non-LTR retrotransposon sequences in both somatic and germline genomes. (**A**) As in Figure 5A. (**B**) As in Figure 5A. Inset shows coverage across the entire contig and position of the retrotransposon gene. **(C**) Alignment of MAC+IES and somatic genomic sequences for Contig_44 retroelement genes from Figure 5A, showing how excision of the central IES deletes part of the endonuclease domain and produces a premature stop codon.

**Figure S6.**
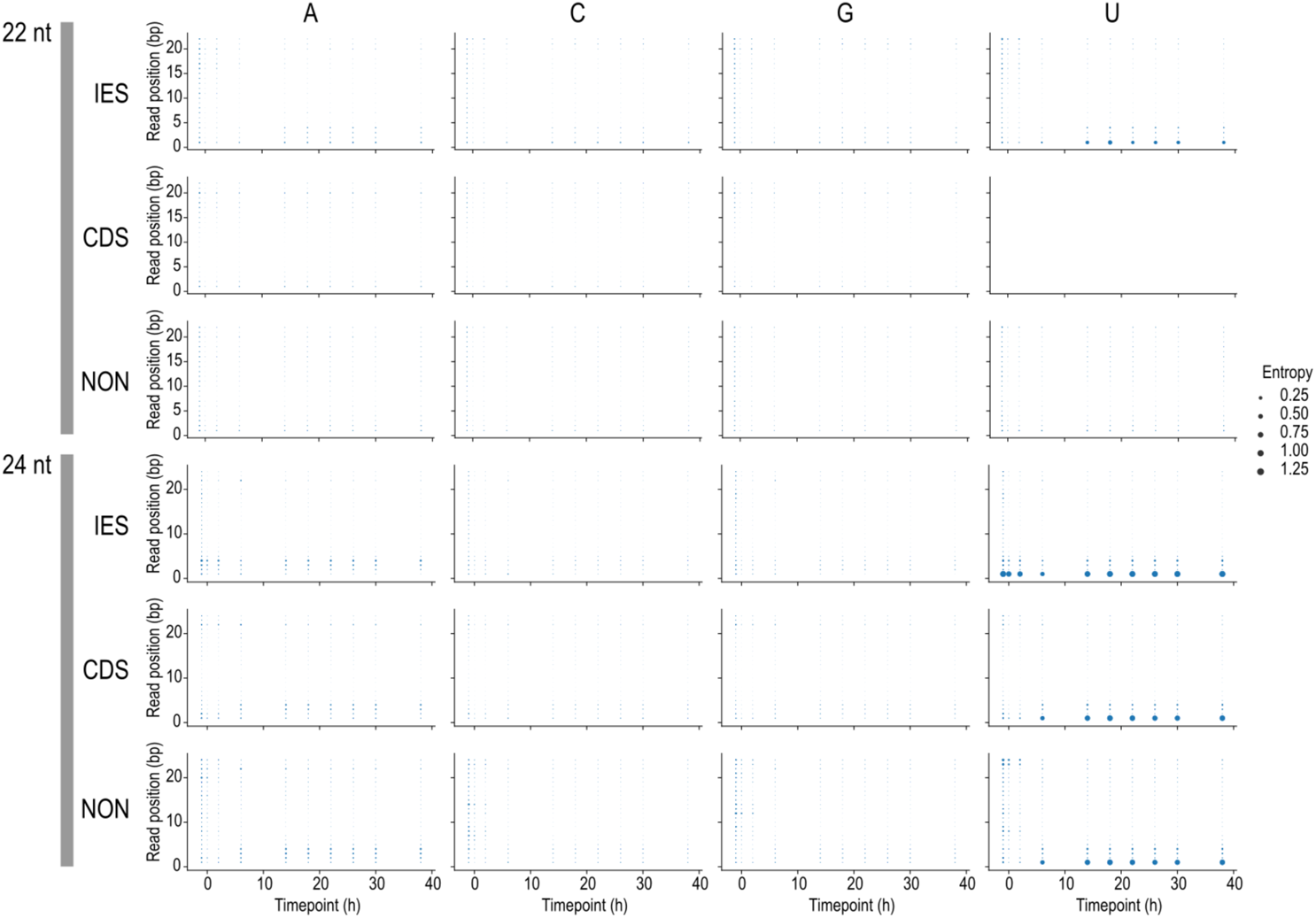
Per-position base entropy of 22 nt and 24 nt sRNAs from developmental time series. Plots show conservation of 5’-U in 24 nt sRNAs. Each plot symbol represents positional sequence entropy (symbol size) for a given nucleotide base (columns) and position in the sRNA sequence (vertical axis) and time point (horizontal axis), in sRNAs mapping to different feature types (rows).

## Methods

General reagents were analytical grade and purchased from Sigma-Aldrich or Merck unless otherwise indicated.

### Ciliate strains origin and cultivation

The strains used and their original isolation localities were: *Blepharisma stoltei* ATCC 30299, Lake Federsee, Germany (Repak, 1968); *Blepharisma stoltei* HT-IV, Aichi prefecture, Japan (Harumoto et al., 1998). Methods for cell cultivation and harvesting of material for sequencing are described in our sister report (Singh et al., 2021).

### Enrichment of micronuclei, isolation and sequencing of genomic DNA

*B. stoltei* ATCC 30299 cells were harvested and cleaned to yield 400 mL of cell suspension (1600 cells/mL). This suspension was twice concentrated by centrifugation (100 g; 2 min; room temperature) in pear-shaped flasks and in 50 mL tubes to ~8 mL. 10 mL chilled Qiagen Buffer C1 (from the Qiagen Genomic DNA Buffer Set, Qiagen no. 19060) and 30 mL chilled, autoclaved deionized water were added. The suspension was mixed by gently inverting the tube until no clumps of cells were visible, and then centrifuged (1300 g; 15 min; 4°C). The pellet was washed with chilled 2 mL Buffer C1 and 6 mL water, mixed by pipetting gently with a wide-bore pipette tip, centrifuged (1300 g; 15 min; 4°C), and resuspended with chilled 2 mL Buffer C1 and 6 mL water by pipetting gently with a wide-bore pipette tip.

The nuclei suspension was layered over a discrete sucrose gradient of 20 mL 10% (w/v) sucrose in TSC medium (0.1% (v/v) Triton X-100, 0.01% (w/v) spermidine trihydrochloride and 5mM CaCl_2_) on top of 40% (w/v) sucrose in TSC medium (Lauth et al., 1976). Gradients were centrifuged (250 g; 10 min; 4°C). 10 to 12 mL fractions were collected by careful pipetting from above, and the nuclei were pelleted by centrifugation (3000 g; 10 min; 4°C). DNA was extracted from pelleted nuclei with the Qiagen Genomic tips 20/G and HMW DNA extraction buffer set (Qiagen no. 19060) according to the manufacturer’s instructions. DNA concentration was measured by the Qubit dsDNA High-Sensitivity assay kit. Fragment size distribution in each sample was assessed by a Femto Pulse analyzer.

*B. stoltei* ATCC 30299 DNA isolated from the MIC-enriched fraction on two separate occasions was used to prepare two sets of DNA sequencing libraries. A low-input PacBio SMRTbell library was prepared without shearing the DNA and was sequenced in the CLR- (continuous long read) sequencing mode on a PacBio Sequel II instrument. Paired-end short-read libraries were prepared for four sucrose gradient fractions (top (T), middle (M), middle lower (ML), bottom (B)) and sequenced with 100 bp BGI-Seq paired-end reads on a BGI-Seq instrument.

### IES prediction from PacBio subreads

PacBio subreads (CLR reads) from a MIC-enriched sample (ENA accession ERR6548140) were aligned to the somatic genome reference assembly (accession PRJEB40285) (Singh et al., 2021) with minimap2 v2.17-r941 (Li, 2018), with options: -ax map-pb --secondary=no -- MD. Mapped reads were sorted and indexed with samtools v1.10 (Li et al., 2009), and then used for predicting IESs with BleTIES MILRAA v0.1.9, with options: --type subreads -- junction_flank 5 --min_ies_length 15 --min_break_coverage 10 -- subreads_pos_max_cluster_dist 5. The BleTIES pipeline has been previously described (Seah and Swart, 2021) and uses spoa v4.0.3 (Vaser et al., 2017) for assembly. After inspecting the initial IES predictions, we removed IES predictions with length <50 bp and retention score <0.075, which we judged to be more likely to be spurious or to have insufficient coverage for an accurate assembly.

Terminal direct repeats (TDRs) at the boundary of a given IES were defined as a sequence of any length that was exactly repeated on both ends of the IES, such that one copy lies within the IES, and the other in the MAC-destined sequence. Because the sequence is identical, it is not possible to determine from sequencing data alone where the physical excision of the IES would occur; such ambiguous excision junctions have been termed “floating IESs” (Sellis et al., 2021). Therefore, TDRs were always reported starting from the left-most coordinate. If the TDR sequence contained 5’-TA-3’, the corresponding IES was also considered to be “TA-bound”, even if the TDR was longer than the 2 bp 5’-TA-3’ sequence.

Reconstructed IES sequences were computationally inserted into the MAC assembly with BleTIES Insert, to produce a hybrid MAC+IES assembly, which approximates the part of the MIC genome that is collinear with the MAC.

### Identification and comparison of IES length classes

Visual inspection of the length distribution of BleTIES-predicted IESs showed sharp peaks every ~10 bp between ~65 and 115 bp. Peak calling on the graph of number of IESs (TA-bound only) vs. length (bp) was performed with the function find_peaks from the Python package scipy.signal v1.3.1 (Virtanen et al., 2020), with height cutoff 100. The ranges for each IES size class were defined with the width at half peak height. In *Paramecium tetraurelia*, where most IESs are TA-bound, the IES termini have a short, weakly conserved inverted repeat (Arnaiz et al., 2012; Klobutcher and Herrick, 1995). To search for similar motifs in *B. stoltei*, sequences flanking TA-bound IES junctions were extracted, with one from each pair reverse-complemented so that the sequences were always in the orientation 5’-(MDS segment)-TA-(IES segment)-3’. Sequence logos of the junctions (10 bp MDS, 14 bp within IES, not including the TA itself) were drawn for each IES length class with Weblogo (Crooks et al., 2004). Only TA-bound IESs were used for the sequence logos because they could be aligned relative to the 5’-TA-3’ repeat, whereas for IESs bound by other types of junctions there is no common reference point to align the boundaries of the IES.

### Probability of a pair of repeated sequences

Under a null model where all bases in a sequence are independently and identically distributed, the probability *P_n_* of having any possible sequence of length *n* bounding a given sequence feature (either a TDR or a TIR) is the sum of probabilities of all possible sequences (each of which notated as *k*) of length *n*, squared: *P_n_* = ∑*_k∊K_p_k_^2^*, which can be transformed to *P_n_* = (∑*_b∊B_p_b_^2^*)*^n^*, where *B* is the alphabet of bases and *p_b_* is the individual probability of each base. The number of possible sequences *k* of length *n* is simply |*K*| = |*B*|*^n^*.

The probability of having a repeat of length at least 2 is equal to the probability of having a repeat of length 2, because all cases of repeat length > 2 implicitly have a repeat of length = 2. Therefore the probability of having a repeat of length exactly *n*, i.e. match in bases 1 to *n*, and mismatch on base *n*+1 is *P_n_* × *Pr* (*mismatch*) = *P_n_* × (1 - ∑*_b∊B_p_b_^2^*). The expected number of TDRs in *Blepharisma* were calculated by using the empirical base frequencies of the MAC+IES genome assembly for *p_b_*, and multiplying this probability by the number of IESs.

### Identification of terminal inverted repeats (TIRs) and palindromes in IESs

The BleTIES-assembled IES sequences for *Blepharisma* were used to identify exact, ungapped terminal inverted repeats (TIRs). Starting from the ends of the IES sequence immediately within the flanking TDRs, each base was compared to the reverse complement of the corresponding base on the opposite end for a match, extending the TIR until a mismatch was encountered, up to a maximum length of 25 bp. The same procedure was used for *Paramecium tetraurelia* using IESs sequences downloaded from ParameciumDB (https://paramecium.i2bc.paris-saclay.fr/files/Paramecium/tetraurelia/51/annotations/ptetraurelia_mac_51_with_ies, accessed 14 October 2021), except that the coordinates of TDRs were first renumbered and extended beyond the “TA” motif if possible, following the BleTIES coordinate numbering convention, in case there are potential TDRs that are longer than a simple TA. The expected number of TIRs of given lengths under a null model was computed as described in “Probability of a pair of sequences”.

Long TIRs (≥10 bp) were clustered by sequence identity to look for IESs of potentially related origin, using the cluster_fast algorithm (Edgar, 2010) implemented in Vsearch v2.13.6 (Rognes et al., 2016) at 80% identity and the CD-HIT definition of sequence identity (-iddef 0). For each resulting cluster of similar TIRs, the cluster centroid was used as the representative sequence shown in Figure TIRS. TDRs associated with each cluster’s IESs were grouped by length, and for each TDR length a degenerate consensus was reported with the degenerate_consensus function of the Bio.motifs module in Biopython v1.74.

Palindromic IESs were defined as IESs that align to their own reverse complement with a sequence identity ≥90% (matching columns over sequence length); this definition was less strict and permitted inexact matches unlike the TIR search, to allow for sequence divergence and assembly errors. IES sequences were aligned with the PairwiseAligner function from Bio.Align in BioPython v1.74, using global mode and parameter match_score = 1.0, with all other scores set to zero.

Palindromic IESs were clustered with Vsearch cluster_fast as described above, except that one sequence (BSTOLATCC_IES35757) was manually removed after inspection of results because it appears to contain two different nested palindromic sequences. Cluster centroids were aligned pairwise as above and used to calculate a matrix of edit distances (matching columns / alignment length). The distance matrix was clustered with average linkage clustering to produce a sequence distance dendrogram with the functions average and dendrogram from scipy.cluster.hierarchy v1.3.1 (Virtanen et al., 2020).

### Comparison of intragenic:intergenic IES ratios

Intragenic vs. intergenic IESs were defined by overlap of predicted IES annotations with “gene” feature annotations on the MAC reference (ENA accession GCA_905310155), using Bedtools v2.30.0 (Quinlan and Hall, 2010) and pybedtools v0.8.1 (Dale et al., 2011).

To test whether the underrepresentation of IESs within gene features was statistically significant, compared to the null hypothesis of IESs and gene feature locations being independently distributed, we assumed that the number of intragenic IESs would follow a binomial distribution with individual probability equal to the fraction of the genome that is covered by gene features. The p-value of the observed number of intragenic IESs would then be equal to the cumulative probability density up to and including the observed value.

### Developmental time series small RNA-seq

Complementary mating strains *B. stoltei* ATCC 30299 and HT-IV were pre-treated with Gamone 2 and Gamone 1 respectively, and then mixed to initiate conjugation as described previously; sRNA and mRNA were isolated from total RNA at the same time points (“Conjugation time course”, (Singh et al., 2021)). sRNA libraries were prepared with the BGISeq-500 Small RNA Library protocol, which selects 18 to 30 nt sRNAs by polyacrylamide gel electrophoresis, and sequenced on a BGISeq 500 instrument.

### Small RNA libraries mapping and comparison

Small RNA libraries were mapped to the MAC+IES assembly with bowtie2 v2.4.2 (Langmead and Salzberg, 2012) using default parameters. Total reads mapping to CDS vs. IES features were counted with featureCounts v2.0.1 (Liao et al., 2014). To account for different total sequence lengths represented by CDSs, IESs, and intergenic regions, the read counts were converted to relative expression values (reads per kbp transcript per million reads mapped, RPKM (Mortazavi et al., 2008)) using the total lengths of each feature type in place of transcript length in the original definition of RPKM, with the following formula:

10^9^ × (reads mapped to feature type) / (total reads mapped × total length of feature type).

Reads mapping to CDSs, IESs, or neither (but excluding tRNA and rRNA features) were extracted with samtools view, with 22 and 24 nt reads extracted to separate files. Read length distributions for each sequence length and feature type were summarized with samtools stats.

### mRNA-seq read mapping

To permit correct mapping of tiny introns RNA-seq data was mapped to the MAC genome using a version of Hisat2 (Kim et al., 2019) with the static variable minIntronLen in hisat2.cpp in the source code lowered to 9 from 20 (https://github.com/Swart-lab/hisat2/; commit hash 86527b9). Hisat2 was run with default parameters and parameters --min-intronlen 9 --max-intronlen 30. It should be noted that spliced-reads do not span introns that are interrupted by an IES due to the low maximum length, however such cases are not expected to occur often.

### Gene prediction and domain annotation

To predict protein-coding genes in IESs, non-IES nucleotides in the MAC+IES assembly were first masked with ‘N’s. The Intronarrator pipeline (https://github.com/Swart-lab/Intronarrator), a wrapper around Augustus (Stanke and Waack, 2003), was run with the same parameters as for the *B. stoltei* MAC genome, i.e. a cut-off of 0.2 for the fraction of spliced reads covering a potential intron, and ≥10 reads to call an intron (Singh et al., 2021). Without masking, gene predictions around IESs were poor, with genuine MDS-limited genes (with high RNA-seq coverage) frequently incorrectly extended into IES regions. The possibility of genes spanning IES boundaries was not catered for.

Domain annotations for diagrams were generated with the InterproScan 5.44-79.0 pipeline (Jones et al., 2014) incorporating HMMER (v3.3, Nov 2019, hmmscan) (Eddy, 2011).

For comparison of transposase-related domain content in MAC vs. MIC, reference sequences were obtained from public databases for *Paramecium tetraurelia* (https://paramecium.i2bc.paris-saclay.fr/files/Paramecium/tetraurelia/51/annotations/ptetraurelia_mac_51_with_ies/), *Tetrahymena thermophila* (http://www.ciliate.org/system/downloads/3-upd-cds-fasta-2021.fasta), and *Oxytricha trifallax* (https://oxy.ciliate.org/common/downloads/oxy/Oxy2020_CDS.fasta, https://knot.math.usf.edu/mds_ies_db/data/gff/oxytri_mic_non_mds.gff). IES gene prediction in *Blepharisma* was hampered by intermittent polynucleotide tract length errors, due to the assembly of IESs from PacBio CLR reads. To mitigate this, a six-frame translation of the MIC-limited genome regions was performed using a custom script, then scanned against the Pfam-A database 32.0 (release 9) (Mistry et al., 2021) with hmmscan (HMMER), with i-E-value cutoff ≤10^-6^. Domains were annotated from the MAC genome with three different methods: using published coding sequences (“cds” in Table S4), six-frame translations (“6ft”), and six-frame translations split on stop codons (“6ft_split”).

### Repeat annotation and clustering

To evaluate the repetitive sequence content in IESs, we applied a repeat prediction and annotation to the combined MAC+IES assembly, instead of clustering whole IESs by sequence similarity. This was so that: (i) Repeats shared between the MDS and IES could be identified. (ii) Complex structures such as nested repeats could be detected. (iii) Repeat families were predicted *de novo*, permitting discovery of novel elements. (iv) Repeats did not have to be strictly identical to be grouped into a family.

Interspersed repeat element families were predicted from the MAC+IES genome assembly with RepeatModeler v2.0.1 (default settings, random number seed 12345) with the following dependencies: rmblast v2.9.0+ (http://www.repeatmasker.org/RMBlast.html), TRF 4.09 (Benson, 1999), RECON (Bao and Eddy, 2002), RepeatScout 1.0.6 (Price et al., 2005), RepeatMasker v4.1.1 (http://www.repeatmasker.org/RMDownload.html). Repeat families were also classified in the pipeline by RepeatClassifier v2.0.1 through comparison against RepeatMasker’s repeat protein database and the Dfam database. Consensus sequences of the predicted repeat families, produced by RepeatModeler, were then used to annotate repeats in the MAC+IES assembly with RepeatMasker, using rmblast as the search engine.

The consensus sequences for rnd-1_family-0 and rnd-1_family-73 were manually curated for downstream analyses. For rnd-1_family-0 (BogoMITE) the original consensus predicted by RepeatModeler for rnd-1_family-0 was 784 bp long, but this was a spurious inverted duplication of the basic ~390 bp unit; the duplication had been favored in the construction of the consensus because RepeatModeler attempts to find the longest possible match to represent each family. For family rnd-1_family-73 (containing BstTc1 transposon), the actual repeat unit was longer than the boundaries predicted by RepeatModeler. In most IESs that contain this repeat (19 of 22), it was flanked by and partially overlapping with short repeat elements from families rnd-4_family-1308 and rnd-1_family-117, which are spurious predictions. Repeat unit boundaries were manually defined by alignment of full length repeats and their flanking regions.

Terminal inverted repeats of selected repeat element families were identified by aligning the consensus sequence from RepeatModeler, and/or selected full-length elements, with their respective reverse complements using MAFFT (Katoh and Standley, 2013) (plugin version distributed with Geneious).

TIRs from the Dfam DNA transposon termini signatures database (v1.1, https://w.dfam.org/releases/dna_termini_1.1/dna_termini_1.1.hmm.gz) (Storer et al., 2021) were searched with hmmsearch (HMMer v3.2.1) against the IES sequences, to identify matches to TIR signatures of major transposon subfamilies.

### Phylogenetic analysis of Tc1/Mariner-superfamily transposases

Repeat family rnd-1_family-1 was initially classified as a “TcMar/Tc2” family transposable element by RepeatClassifier. 30 full length copies (>95% of the consensus length) were annotated by RepeatMasker, all of which fell within IESs and contained CDS predictions. However, CDSs were of varying lengths because of frameshifts caused by indels, which may be biological or due to assembly error; nonetheless, the nucleotide sequences had high pairwise identity (about 98%, except for one outlier). We chose BSTOLATCC_MIC4025 as the representative CDS sequence for phylogenetic analysis because it was one of the longest predicted and both predicted Pfam domains (HTH_Tnp_Tc5 and DDE_1) appeared to be intact.

For repeat family rnd-1_family-73, the initial classification was “DNA/TcMar-Tc1”. As described above, CDS predictions were of variable lengths, and the longest CDSs were not necessarily the best versions of the sequence because of potential frameshift errors. For phylogenetic analysis, we chose BSTOLATCC_MIC48344 as the representative copy, because a complete *DDE_3* Pfam domain was predicted by HMMER that could align with other DDE/D domains from reference alignments described below.

The representative CDSs of the rnd-1_family-1 and rnd-1_family-73 transposases were aligned with MAFFT (E-INS-i mode) against a published DDE/D domain reference alignment (Supporting Information Dataset_S01 of (Yuan and Wessler, 2011)) to identify the residues at the conserved catalytic triad and the amino acid distance between the conserved residues.

For the phylogenetic analysis of the DDE/D domains in the Tc1/Mariner superfamily, both MAC- and MIC-limited genes containing DDE_1 and DDE_3 domains were separately aligned for each Pfam domain with MAFFT v7.450 (algorithm: E-INS-i, scoring matrix: BLOSUM62, Gap open penalty: 1.53) and trimmed to the DDE/D domain with Geneious and incomplete domains were removed. As reference, 204 sequences from a published alignment (Additional File 4 of (Dupeyron et al., 2020)) were selected to represent the 53 groups defined in that study, choosing only complete domains (with all three conserved catalytic residues) and all *Oxytricha trifallax* TBE and *Euplotes crassus* Tec transposase sequences. Thirteen *Paramecium* Tc1/Mariner DDE/D domain consensus sequences were added (Additional File 4 of (Guérin et al., 2017)). Sequences were aligned with MAFFT (E-INS-i mode) and trimmed to only the DDE/D domain boundaries with Geneious. Phylogeny was inferred with FastTree2 v2.1.11 (Price et al., 2010) using the WAG substitution model. The tree was visualized with Dendroscope v3.5.10 (Huson and Scornavacca, 2012), rooted with bacterial IS630 sequences as outgroup

### Phylogenetic analysis of retrotransposon-derived sequences

All the nucleotide sequences ≥500 bp for the repeat families identified by RepeatClassifier as LINE or LINE/RTE-x: rnd-1_family-273, rnd-1_family-276 and rnd-4_family-193 were aligned to one another with MAFFT v7.450 (automatic algorithm) (Katoh and Standley, 2013), with the option to automatically determine sequence direction (via the MAFFT plugin for Geneious Prime (Kearse et al., 2012)). Since the alignment appeared to be poor between the rnd-4-family-193 sequences and the rest, we generated separate alignments for this family from the other two, also with MAFFT (E-INS-i mode). Maximum likelihood phylogenies were generated by PhyML (Guindon et al., 2010) version 3.3.20180621 with the HKY85 substitution model.

## Data availability

Annotated draft MAC+IES genome for *Blepharisma stoltei* strain ATCC 30299 (European Nucleotide Archive (ENA) Bioproject PRJEB46944 under accession GCA_914767885). IES sequences and annotations, MAC gene predictions with intervening IESs, and gene predictions within IESs (EDMOND, doi:<u>10.17617/3.83</u>; genome browser, https://bleph.ciliate.org. Sequencing data for the MIC-enriched nuclear fractions (PacBio CLR reads: ENA accession ERR6510520 and ERR6548140; BGI-seq reads: ENA accessions ERR6474675, ERR6496962, ERR6497067, ERR6501836). Small RNA libraries from developmental time series (ENA Bioproject PRJEB47200 under accessions ERR6565537-ERR6565561). Repeat family predictions and annotations by RepeatModeler and RepeatMasker (EDMOND, doi:<u>10.17617/3.82</u>). Alignment and phylogeny of Tc1/Mariner superfamily transposase domains (EDMOND, doi:<u>10.17617/3.JLWBFM</u>)

## Abbreviations

IES: internally eliminated sequence
LTR: long terminal repeat
MAC: macronucleus
MIC: micronucleus
MITE: miniature inverted-repeat transposable element
MITIES: miniature inverted-repeat transposable internally eliminated sequences
TDR: terminal direct repeat
TIR: terminal inverted repeat
TSD: target site duplication

## Acknowledgements

We thank C. Lanz for assistance with DNA quality control, A. Noll for computer system administration, L.A. Klobutcher, G. Herrick, O. Weichenrieder and A. Streit for helpful discussions. Research reported in this publication was supported by the National Institutes of Health (award No. P40OD010964) to N.A.S and the Max Planck Society.

## Author contributions

Data curation: B.K.B.S., E.C.S., M.S.1. Formal analysis: B.K.B.S., M.S.1, E.C.S., C.W. Funding acquisition: N.S., E.C.S. Investigation: B.K.B.S., M.S.1., E.C.S. Methodology: B.K.B.S., M.S.1., E.C.S., M.S.2, T.H., A.S., C.E. Resources: M.S.2., T.H., C.W., B.H., A.B., N.S. Software: B.K.B.S., E.C.S., M.S.1, A.B., N.S. Supervision: M.S.2, T.H., E.C.S. Visualization: B.K.B.S., M.S.1., E.C.S. Writing – original draft: B.K.B.S., M.S.1., E.C.S. Writing – review & editing: B.K.B.S., M.S.1, E.C.S., M.S.2, T.H., A.S., C.W., N.S.

