## Supplemental Information for "MITE infestation of germline accommodated by genome editing in *Blepharisma*"

### Supplemental Results

#### IES assembly from short reads vs. long reads

IESs predicted from short read sequencing (ParTIES prediction) had higher retention scores than those from the long read library (BleTIES prediction) (Figure S1A). However, IESs of size ~390 bp were largely absent from the ParTIES prediction, despite being an abundant size class in the BleTIES prediction (Figure S1A). We attributed this to the repetitive sequence content in the ~390 bp IESs, which contained a highly conserved repeat element. ParTIES pools all putative IES-containing sequences in the whole library for reassembly before aligning the resulting contigs to the MAC to identify IESs [(Denby Wilkes et al., 2016)](https://sciwheel.com/work/citation?ids=4034499&pre=&suf=&sa=0), whereas BleTIES performs a separate targeted assembly for each IES. IESs which contain repetitive sequence content would be less likely to be accurately predicted by ParTIES because reads originating from different IESs would be assembled together into a hybrid contig that cannot be aligned to the MAC reference. Therefore, we used the BleTIES-predicted IESs for all subsequent analyses.

#### MIC sequence coverage and telomeric content

Per-IES retention scores from BleTIES had a median of 0.195, indicating that about 20% of the read library originated from the MIC genome (Figure S1A). The average coverage of IES-containing reads underlying the predicted IESs was ~45x, but given the distribution of coverage values, we expect that more IESs could be assembled with greater sequencing coverage.

Reads originating from MIC (IES-containing) also contained less telomeric sequence (0.0228% of total read length) than MAC reads (IES-lacking) (2.98%). This was consistent with our previous observation that the MAC genome was fragmented into telomere-bound minichromosomes of ~130 kbp length, presumably, like other ciliates, from longer and more contiguous MIC precursor chromosomes [(Singh et al., 2021)](https://sciwheel.com/work/citation?ids=12158331&pre=&suf=&sa=0).

#### Periodic IES length distribution

*Paramecium* and *Blepharisma* IESs differ in the following ways: (i) *Paramecium* IESs are shorter on average, with a first peak at ~27 bp compared to ~65 bp in *Blepharisma*; (ii) *Paramecium* has a “missing” second peak at ~36 bp; and (iii) the first peak (27 bp) is the highest in *Paramecium* and heights decrease thereafter (except for the “missing” peak), whereas in *Blepharisma* the second (72 bp) and sixth (110 bp) peaks are the highest. Geometric constraints of the excisase complex bound to DNA have been proposed as an explanation for the periodic length distribution and missing second peak of *Paramecium* [(Arnaiz et al., 2012)](https://sciwheel.com/work/citation?ids=2201947&pre=&suf=&sa=0). In this model, the shortest ~27 bp *Paramecium* IESs (peak 1) represent the length of DNA required to bridge the cleavage sites on two subunits of the excisase, whereas peaks 3 and above represent DNA with an intervening loop, and the ~10 bp length periodicity corresponds to the ~10 bp period of the dsDNA helix. Peak 2 is “forbidden” because it is too long for the active excisase complex but too short to form a loop. The periodic IES lengths in *Blepharisma* can also be explained by this model because the last periodic peak (110 bp) is still below the persistence length of DNA, however they are not as short as those in *Paramecium*. This suggests that all the periodic IESs *Blepharisma* IESs are also looped, but that their excisase is unable to operate on the very short loops that may occur in *Paramecium* (down to 44 bp).

The secondary maximum peak at 110 bp may represent a historical wave of IES proliferation: Assuming that new IESs usually start out longer than 115 bp, they gradually decay in length with time. Once they reach the periodic length range, they are “captured” in optimal excision peaks lengths, and will eventually accumulate at the shortest-length peak (72 bp in *Blepharisma*, 27 bp in *Paramecium*), which represent the most abundant size class. Therefore, the uppermost periodic peak (110 bp) may contain IESs that proliferated sufficiently long ago that the IESs have been degraded to ~110 bp length, but not long enough to filter down to the shorter length peaks. Alternatively, it could represent a secondary conformational optimum for the excisase complex, in addition to the primary optimum at 72 bp.

#### Additional palindromic IESs (putative MITE IESs)

Of the 376 palindromic IESs (≥90% identity in self-alignment) identified, 153 (40.7%) fell within the ~228 bp IES length peak observed before (225-231 bp, Table S1), although some palindromic IESs were within the periodic IES length range (Figure S2). However, when clustered at 90% sequence identity, palindromic IESs in the ~228 bp length range actually fell into several clusters, suggesting that this peak was composed of several families of palindromic IESs which happened to have a similar length, rather than a single family. This was confirmed by pairwise distances and visual inspection of the multiple sequence alignment of the cluster centroids (Figure S2B, S2C). Some clusters had recognizable homology to each other, but many were over 40% divergent.

#### “Cryptic” IESs in the MAC genome

In addition to conventional IESs, “cryptic” IES excision can occasionally occur, which is the low frequency excision of MDS sequences that are incorrectly recognized as an IES by the excision machinery [(Duret et al., 2008; Swart et al., 2014)](https://sciwheel.com/work/citation?ids=4034486,326423&pre=&pre=&suf=&suf=&sa=0,0). Cryptic IESs were identified by mapping MAC reads back onto the MAC reference assembly and looking for pileups of deletions relative to the reference.

10048 potential cryptic IESs ≥50 bp were detected, of which 5635 (56.1%) were TA-bound, and 1328 (13.2%) bound by other TDRs. The fraction of TA-bound cryptic IESs was lower than that of conventional IESs, which could be partially attributed to misprediction of cryptic IESs, because a lower coverage threshold was used to detect them compared to conventional IESs. IES-negative forms represent only a minority of reads at cryptic IES positions (median retention score 95%, i.e. only 5% of reads are IES-negative, Figure S1D). Conversely, true IESs appear to be efficiently excised from MAC DNA (Figure S1B). Nonetheless, the length threshold of cryptic IESs has a clear peak at ~72 bp, corresponding to the most abundant periodic IES size class found previously, and less prominent peaks at other size classes (82, 92, 101, 110). Furthermore, the fraction of cryptic IESs in the ~72 bp size class that were TA-bound was higher than average, at 66.0%. The sequence logo of TA-bound cryptic IES junctions did not show any obvious sequence bias apart from a T/A immediately after the “TA”, but the sequence logo for only the ~72 bp cryptic IESs shows a slight TTT bias from position 6 after the “TA” (Figure S1F), which resembles the motif found in conventional IESs (Figure 1D).

The most common TDR sequence of cryptic IESs was “TA”. Simple alternations of T/A were also common, as well as sequences containing “TTA” or “TAA”. Unlike the conventional IESs, cryptic IESs with “TAA” or “TTA” TDRs did not form a distinct size class at ~390 bp corresponding to the BogoMITEs, but were distributed similarly to the other cryptic IESs, with a peak at 72 bp (Figure S1E). Therefore, it is likely that TTA/TAA could represent an intrinsic cut site preference of the domesticated excisase (or one of them).

#### Intragenic:intergenic ratio for different IES size classes

We also compared the ratios of intragenic to intergenic IESs for different IES size classes. We hypothesized that IESs belonging to the different ranges of IES size classes (periodic vs. non-periodic) may not have the same excision efficiency and would hence experience different selective regimes. For example, an intragenic IES with poor excision efficiency would be more negatively selected against than one with better efficiency. Compared to the overall intragenic:intergenic ratio of 2.33 (i.e. 70% intragenic), the IESs that belonged to the most abundant size class (~72 bp) were more likely to be intragenic (ratio 2.70, 73% intragenic, p < 0.001) than IESs as a whole (Table S6). Other size classes also had higher or lower ratios compared to the expectation but they were not statistically significant with our relatively conservative p-value cutoff.

#### Catalytic triad in DDE/D-superfamily transposases

For the each of the families in DDE/D superfamily, we observed one of eight PiggyBac domains, four of a total of nine DDE_1/DDE_3 domains, three of five DDE_Tnp_IS1595 domains and five of six instances of the MULE domains in the MAC genome, with intact catalytic triads. The presence of the catalytic triad in the MIC instances of these domains was more varied. None of the PiggyBac domains had a complete catalytic triad, though the longest cORF contained an almost complete catalytic triad, where the second Aspartate residue appeared to be translocated by one amino acid. For the DDE1/DDE_3 domain-carrying MIC protein, only fifteen of the forty seven lacked the complete triad. Three of five MIC-limited DDE_Tnp_IS1595 domains and nine of ten MIC-limited MULE domains lacked the catalytic triad.

#### Sequence diversity of MAC-limited retrotransposon-derived repeats

Repeat families rnd-1_family-276 and rnd-1_family-273 defined by RepeatModeler/ RepeatClassifier had partially overlapping membership, but largely correspond to two clusters of related sequences. Between clusters, there was 60 to 69% pairwise nucleotide identity (Figure 5A), but within clusters, sequence identities were very high (>97%). For example, in addition to the seven high-identity, ~4.1 kbp long copies of repeat daily rnd-1_family-276 (Main Text), seven shorter sequences with high identity to these (>97%) were found at other genomic locations (five in the cruft MAC+IES region). Three > 3 kb sequences present on different MAC genome contigs are > 98% identity at the nucleotide level with additional high identity copies present in the “cruft” genome portion.

An additional retrotransposon-derived repeat family, rnd-4_family-193 (Table S5; Figure 5B) was more distantly related (28-35% nucleotide identity relative to long sequences from the other two families) and more divergent within the family itself. Among the rnd-4_family-193 sequences classified, only one copy was relatively long (4.6 kbp), and no sequences showed additional long, high-identity matches as was observed for rnd-1_family-273 and rnd-1_family-276. The next longest rnd-1_family-193 sequences were ~2.0 kbp, and thus too short to encode a complete retrotransposase with both endonuclease and reverse transcriptase domains.

#### Expression of non-LTR-retrotransposon-derived sequences

In *Tetrahymena* cells, retrotransposon transcription was below the detection limits in vegetatively growing and starved cells, first observed when meiosis occurs, disappearing with time [(Fillingham et al., 2004)](https://sciwheel.com/work/citation?ids=2201960&pre=&suf=&sa=0). In *Oxytricha* expression of LINE retroelements is prominent well after meiosis and negligible prior to this [(Chen et al., 2014)](https://sciwheel.com/work/citation?ids=372905&pre=&suf=&sa=0). In contrast, the expression of the *Blepharisma* RVT_1 genes was negligible in starved cells and throughout development [(Singh et al., 2021)](https://sciwheel.com/work/citation?ids=12158331&pre=&suf=&sa=0). Such barely detectable expression throughout development was not observed for any of the previously proposed putative domesticated transposase families in the *B. stoltei* MAC genome [(Singh et al., 2021)](https://sciwheel.com/work/citation?ids=12158331&pre=&suf=&sa=0). However, none of *Blepharisma*’s putative domesticated transposases are anywhere near as abundant as the retrotransposon repeats in the MAC genome, let alone show signs of substantial recent replication.

#### Rate of development post-conjugation

The timing of sRNA expression and turnover in *Blepharisma* appeared to be slower than in a similar experiment in *Tetrahymena*, hence the earlier timepoints of our *Blepharisma* experiment captured intermediate stages not observed in *Tetrahymena*. At 6, 14 h timepoints in *Blepharisma*, MDSs have comparable 24 nt sRNA coverage to IES regions, whereas by 3 h after mixing of complementary mating types in *Tetrahymena*, about 80% of scnRNAs mapped to IESs [(Schoeberl et al., 2012)](https://sciwheel.com/work/citation?ids=2310080&pre=&suf=&sa=0). It is likely that development generally proceeds faster in *Tetrahymena*.

### Supplemental Methods

#### IES prediction from BGIseq short reads

BGIseq reads (100 bp, paired end) from MIC-enriched sample “AT10” (ENA accession ERR6501836) were mapped to the MAC reference assembly with bowtie2 v2.4.2 on local mode within the ParTIES pipeline [(Denby Wilkes et al., 2016)](https://sciwheel.com/work/citation?ids=4034499&pre=&suf=&sa=0). We modified ParTIES (based on v1.02) to use SPAdes v3.15.2 [(Prjibelski et al., 2020)](https://sciwheel.com/work/citation?ids=9670085&pre=&suf=&sa=0) instead of Velvet [(Zerbino and Birney, 2008)](https://sciwheel.com/work/citation?ids=162676&pre=&suf=&sa=0) to assemble IES+ sequences (<https://github.com/Swart-lab/ParTIES>, branch “custom” commit f04ad7e2), since Velvet kept crashing on our data.

#### MIC read binning and telomere annotation

Internally error-corrected circular consensus sequence (CCS) reads were generated from the above CLR library with PacBio ccs v4.2.0 (<https://github.com/PacificBiosciences/ccs>), with 26.1% of ZMWs generating CCSs. CCS reads were mapped to the MAC reference assembly with minimap2 with the same parameters as CLR reads, except for option -ax asm20. The mapping and IES annotation were used to calculate per-read IES retention scores. Reads were binned with the MILCOR module of BleTIES into putative MAC (score <0.1) and putative MIC (score >0.9). Total telomere sequence length in the binned MAC and MIC reads were calculated with a Python regular expression search for the telomere repeat 5’-CCCTAACA-3’ and its reverse complement.

#### Accommodation of IESs in annotation feature tables

IESs in the MAC+IES genome assembly submitted to ENA were annotated as “iDNA” features, which was the closest-fitting feature type supported by INSDC feature tables (<https://www.insdc.org/files/feature_table.html>), although IESs are included in the Sequence Ontology (http://www.sequenceontology.org/browser/current_release/term/SO:0000671).

We also note that IES features can potentially create confusion for certain applications when an IES is present within a CDS feature. This is because in GFF3 and INSDC feature tables, introns are usually defined implicitly when a CDS is split into multiple segments, rather than being separately annotated. Therefore with a MAC+IES genome assembly and gene annotations from public sequence databases like ENA, one has to be careful to check whether the CDS is interrupted by introns or IESs, or both.

#### IES retention scores in MAC enrichment library

PacBio HiFi reads from a MAC enrichment library were mapped to the *B. stoltei* MAC reference assembly with minimap2, sorted and indexed with samtools. Retention scores for previously predicted IESs in the MAC were calculated with BleTIES MILRET with default parameters.

#### Annotation of cryptic IESs

PacBio HiFi reads representing *B. stoltei* ATCC 30299 MAC DNA (ENA ERR5873334, ERR5873783) were mapped onto the MAC reference (GCA_905310155) with minimap2, and sorted and indexed with samtools as described above. The mapping was processed with BleTIES MILRAA with options --type ccs --fuzzy_ies --min_ies_length 15 --min_break_coverage 2 --min_del_coverage 2. IES predictions that overlapped with telomere regions and “cruft” contigs were removed. Those IES predictions that represented deletions relative to the reference assembly were considered to represent possible cryptic IESs. TDRs and sequence logos of cryptic IES junctions were defined and drawn as described above for conventional IESs.

#### Intragenic:intergenic IES ratios for specific IES size classes

We tested whether specific IES length classes were more or less depleted within gene features, compared to all IESs as a whole (two-tailed test, null hypothesis: all IES length classes have equal probability of being intragenic). The intragenic vs. intergenic membership of IESs was held constant, but their assigned lengths were randomly permuted (without replacement) to obtain 1000 pseudoreplicates. For each of the 10 IES size classes defined in Table S1, the p-value was calculated as the fraction of pseudoreplicates where the simulated number of intragenic IESs was more than the actual observed value. The uncorrected p-value threshold (0.05) was adjusted to 0.005 after applying a Bonferroni correction for 10 tests, yielding <0.0025 and >0.9975 as the two-tailed p-value thresholds.

#### Sequence logos for Bogo and BogoMITE repeat boundaries

To generate sequence logos for the Bogo and BogoMITE repeat boundaries, full length sequences for repeat families rnd-1_family-1 (>1800 bp) and rnd-1_family-0 (between 385-395 bp) and their flanking 10 bp were extracted and reverse complemented if necessary to be in the same orientation. Both repeats have a poly-C run in the TIR whose variable length results in misalignment at the repeat boundaries. Therefore, to ensure that the TSDs are properly aligned, the TSD and first three bases of the TIR on each boundary flanking the repeats were identified with a regular expression ..ACTC or its reverse complement GAGT..; the sequences within those boundaries were aligned with the E-INS-i algorithm from MAFFT v7.475. Alignment columns comprising >90% gaps were removed. The degapped alignment was concatenated with the flanking TSD+TIR removed earlier, and then used to generate sequence logos of the repeat boundary regions with Weblogo v3.7.5.
