## Supplementary material for "MITE infestation of germline accommodated by genome editing in *Blepharisma*": Table S1

IES size classes, defined by peak calling on the length distribution of TA-bound IESs. Lower and upper lengths per size class are inclusive. Only TA-bound IESs on main assembly contigs are included in the counts and total lengths.

| Peak center (bp) | Lower bound (bp) | Upper bound (bp) | No. IESs | Total IES length (bp) |
|---|---|---|---|---|
| 65 | 64 | 66 | 207 | 13415 |
| 72 | 70 | 74 | 2717 | 195724 |
| 82 | 80 | 84 | 1035 | 85017 |
| 92 | 90 | 94 | 819 | 75458 |
| 101 | 99 | 103 | 688 | 69638 |
| 110 | 108 | 112 | 1592 | 175060 |
| 153 | 151 | 155 | 336 | 51478 |
| 174 | 173 | 175 | 377 | 65575 |
| 228 | 225 | 231 | 769 | 175422 |
| 389 | 385 | 393 | 876 | 340798 |
