## Supplementary material for "MITE infestation of germline accommodated by genome editing in *Blepharisma*": Table S2

Summary of RepeatMasker annotations in *B. stoltei* MAC+IES assembly for each repeat class, as classified by RepeatClassifier. The most abundant repeat family (rnd-1_family-0) is also listed separately, despite being unclassified. Only one family, rnd-1_family-1, is classified as DNA/TcMar-Tc2. Total annotated length does not account for overlapping annotations.

| Class | Number of annotated elements | Total sequence length annotated (bp) |
|---|---|---|
| Unknown (excluding rnd-1_family-0) | 41836 | 11279760 |
| rnd-1_family-0 (Unknown) | 8369 | 2692873 |
| Simple_repeat | 6878 | 613736 |
| Low_complexity | 2511 | 123672 |
| rnd-1_family-1 (DNA/TcMar-Tc2) | 539 | 104263 |
| LINE/RTE-X | 94 | 51679 |
| LTR/Pao | 39 | 10475 |
| LINE | 28 | 38630 |
| DNA/TcMar-Tc1 | 24 | 38070 |
| Unknown/Helitron-2 | 23 | 11025 |
