## Supplementary material for "MITE infestation of germline accommodated by genome editing in *Blepharisma*": Table S3

Top five most abundant repeat families in specific IES size classes (defined in Table S1). Repeats comprising > 20% of the total IES length of particular size classes are highlighted in bold font.

| Repeat family | Number | Fraction of total IES length | IES size class (peak center bp) |
|---|---|---|---|
| rnd-1_family-397 | 712 | 0.044433 | 65 |
| A-rich | 134 | 0.008362 | 65 |
| rnd-1_family-157 | 67 | 0.004181 | 65 |
| rnd-1_family-151 | 65 | 0.004056 | 65 |
| rnd-4_family-596 | 64 | 0.003994 | 65 |
| A-rich | 2735 | 0.012229 | 72 |
| rnd-1_family-438 | 1898 | 0.008487 | 72 |
| rnd-1_family-397 | 1511 | 0.006756 | 72 |
| rnd-1_family-398 | 508 | 0.002271 | 72 |
| (T)n | 450 | 0.002012 | 72 |
| A-rich | 741 | 0.007556 | 82 |
| rnd-1_family-397 | 441 | 0.004497 | 82 |
| rnd-2_family-94 | 182 | 0.001856 | 82 |
| rnd-1_family-0 | 171 | 0.001744 | 82 |
| (AT)n | 153 | 0.001560 | 82 |
| A-rich | 570 | 0.006505 | 92 |
| rnd-1_family-205 | 400 | 0.004565 | 92 |
| rnd-1_family-0 | 270 | 0.003081 | 92 |
| rnd-2_family-11 | 209 | 0.002385 | 92 |
| rnd-3_family-853 | 160 | 0.001826 | 92 |
| A-rich | 801 | 0.009991 | 101 |
| rnd-3_family-853 | 679 | 0.008469 | 101 |

| | | | |
|---|---|---|---|
| rnd-4_family-1308 | 277 | 0.003455 | 101 |
| rnd-1_family-0 | 174 | 0.002170 | 101 |
| (TATAA)n | 128 | 0.001596 | 101 |
| rnd-3_family-853 | 2275 | 0.011457 | 110 |
| A-rich | 1134 | 0.005711 | 110 |
| rnd-2_family-11 | 336 | 0.001692 | 110 |
| rnd-2_family-94 | 247 | 0.001244 | 110 |
| rnd-1_family-210 | 239 | 0.001204 | 110 |
| **rnd-1_family-87** | **14889** | **0.236551** | **153** |
| rnd-1_family-203 | 5865 | 0.093181 | 153 |
| rnd-1_family-181 | 1331 | 0.021146 | 153 |
| rnd-1_family-93 | 1059 | 0.016825 | 153 |
| rnd-4_family-669 | 621 | 0.009866 | 153 |
| **rnd-1_family-82** | **19793** | **0.268358** | **174** |
| rnd-1_family-80 | 6065 | 0.082231 | 174 |
| rnd-1_family-93 | 4335 | 0.058775 | 174 |
| rnd-1_family-65 | 3970 | 0.053826 | 174 |
| rnd-1_family-224 | 2951 | 0.040010 | 174 |
| rnd-1_family-160 | 18889 | 0.093361 | 228 |
| rnd-1_family-10 | 10109 | 0.049965 | 228 |
| rnd-1_family-16 | 9541 | 0.047158 | 228 |
| rnd-1_family-14 | 9344 | 0.046184 | 228 |
| rnd-4_family-669 | 9054 | 0.044750 | 228 |
| **rnd-1_family-0** | **294091** | **0.684765** | **389** |
| rnd-4_family-95 | 38821 | 0.090391 | 389 |
| rnd-3_family-190 | 9012 | 0.020984 | 389 |
| rnd-4_family-1308 | 5945 | 0.013842 | 389 |
| rnd-1_family-25 | 4430 | 0.010315 | 389 |
