## Supplementary material for "MITE infestation of germline accommodated by genome editing in *Blepharisma*": Table S4

Numbers of transposase-related Pfam domains in MAC vs. MIC-limited sequences (IESs) for different ciliate species, based on hmmscan search of six-frame translations (6ft), six-frame translations split on stop codons (6ft split, shown in Figure 4E), or predicted coding sequences only (cds).

| | *Blepharisma stoltei* | | | | *Paramecium tetraurelia* | | | | *Tetrahymena thermophila* | | | | *Oxytricha trifallax* | | | |
|---|---|---|---|---|---|---|---|---|---|---|---|---|---|---|---|---|
| | MAC | | | MIC | MAC | | | MIC | MAC | | | MIC | MAC | | | MIC |
| Domain | 6ft split | 6ft | cds | ies 6ft | 6ft split | 6ft | cds | ies 6ft | 6ft split | 6ft | cds | ies 6ft | 6ft split | 6ft | cds | ies 6ft |
| DDE_1 | 1 | 0 | 3 | 14 | 0 | 0 | 0 | 0 | 0 | 0 | 1 | 83 | 0 | 0 | 0 | 0 |
| DDE_3 | 2 | 2 | 6 | 15 | 1 | 3 | 0 | 7 | 0 | 0 | 3 | 868 | 0 | 0 | 2 | 451 |
| DDE_Tnp_1_7 | 7 | 0 | 9 | 5 | 1 | 0 | 9 | 0 | 3 | 0 | 3 | 42 | 0 | 0 | 0 | 0 |
| DDE_Tnp_IS1595 | 2 | 3 | 5 | 3 | 0 | 0 | 0 | 0 | 0 | 0 | 1 | 138 | 1 | 7 | 7 | 28 |
| Exo_endo_phos_2 | 5 | 0 | 12 | 1 | 0 | 0 | 1 | 0 | 0 | 0 | 1 | 5 | 0 | 0 | 0 | 1 |
| HTH_Tnp_Tc5 | 1 | 1 | 4 | 22 | 5 | 9 | 12 | 3 | 0 | 1 | 1 | 56 | 0 | 1 | 2 | 1 |
| MULE | 3 | 2 | 6 | 7 | 0 | 0 | 0 | 0 | 0 | 0 | 0 | 2 | 2 | 8 | 6 | 0 |
| RVT_1 | 10 | 4 | 27 | 8 | 0 | 0 | 3 | 4 | 0 | 0 | 3 | 38 | 0 | 0 | 0 | 45 |
| Transposase_mut | 0 | 0 | 0 | 1 | 0 | 0 | 0 | 0 | 0 | 0 | 0 | 0 | 2 | 0 | 1 | 0 |
| Dimer_Tnp_hAT | 0 | 0 | 0 | 0 | 0 | 0 | 0 | 0 | 0 | 3 | 1 | 136 | 0 | 0 | 0 | 0 |
| HTH_Tnp_1 | 0 | 0 | 1 | 0 | 0 | 0 | 0 | 0 | 0 | 0 | 0 | 2 | 0 | 0 | 0 | 0 |
| HTH_Tnp_Tc3_2 | 0 | 0 | 0 | 0 | 0 | 0 | 0 | 0 | 0 | 0 | 0 | 124 | 0 | 0 | 0 | 0 |
| Transposase_1 | 0 | 0 | 0 | 0 | 0 | 0 | 0 | 0 | 0 | 0 | 0 | 30 | 0 | 0 | 0 | 0 |
| Helitron_like_N | 0 | 0 | 0 | 0 | 0 | 0 | 0 | 0 | 0 | 0 | 0 | 0 | 0 | 0 | 0 | 3 |
| Tnp_zf-ribbon_2 | 0 | 0 | 0 | 0 | 0 | 0 | 0 | 0 | 0 | 0 | 1 | 0 | 0 | 0 | 0 | 0 |
| DDE_Tnp_1 | 0 | 0 | 0 | 0 | 0 | 0 | 0 | 0 | 0 | 0 | 0 | 0 | 0 | 3 | 2 | 0 |
| DDE_Tnp_1_2 | 0 | 0 | 0 | 0 | 0 | 0 | 0 | 0 | 0 | 0 | 0 | 0 | 0 | 1 | 1 | 0 |

| | | | | | | | | | | | | | | | | |
|---|---|---|---|---|---|---|---|---|---|---|---|---|---|---|---|---|
| DDE_Tnp_1_3 | 0 | 0 | 0 | 0 | 0 | 0 | 0 | 0 | 0 | 0 | 0 | 0 | 0 | 0 | 1 | 0 |
| DDE_5 | 0 | 0 | 0 | 0 | 0 | 0 | 0 | 0 | 0 | 0 | 0 | 0 | 0 | 2 | 0 | 0 |
