## Supplementary material for "MITE infestation of germline accommodated by genome editing in *Blepharisma*": Table S5

Summary of RepeatMasker annotations for individual repeat families that were classified by RepeatClassifier. Repeats identified predominantly in IESs are highlighted in bold. RepeatClassifier classifications that appear to be errors or spurious annotations are surrounded in parentheses: family rnd-1_family-283 mostly comprises ubiquitin sequences, whereas rnd-4_family-1389 contains abundant WD40 repeats.

| | | | All copies | | | | Full length copies only | | | |
|---|---|---|---|---|---|---|---|---|---|---|
| **Repeat family** | **Class (RepeatClassifier)** | **Cons len. (bp)** | **No.** | **Median copy len. (bp)** | **Total len. (bp)** | **No. on IESs** | **No.** | **Total len. (bp)** | **No. on IESs** | **% div. vs. cons** |
| **rnd-1_family-1** | **TcMar/Tc2** | **1833** | **539** | **91** | **104802** | **505** | **30** | **54844** | **30** | **0.5** |
| **rnd-1_family-73** | **DNA/TcMar-Tc1** | **1949** | **28** | **1640** | **38098** | **27** | **22** | **36273** | **22** | **0.6** |
| rnd-1_family-273 | LINE | 3618 | 23 | 1319 | 38653 | 2 | 6 | 21708 | 0 | 16.9 |
| rnd-1_family-276 | LINE/RTE-X | 3270 | 15 | 723 | 16197 | 4 | 2 | 6451 | 1 | 2.95 |
| rnd-1_family-283 | (LTR/Pao) | 358 | 39 | 339 | 10514 | 3 | 24 | 8438 | 0 | 16.25 |
| rnd-4_family-193 | LINE/RTE-X | 4628 | 79 | 279 | 35576 | 36 | 1 | 4628 | 1 | 9.5 |
| rnd-4_family-1389 | (Unknown/Helitron-2) | 2108 | 24 | 268 | 11049 | 0 | 1 | 2108 | 0 | 5.8 |
