## Supplementary material for "MITE infestation of germline accommodated by genome editing in *Blepharisma*": Table S6

Counts of intra- vs. intergenic localization for IESs in different size classes (defined in Table S1).

| IES size class (peak center bp) | Intergenic | Intragenic | IES size class type | Ratio intra:inter - genic | Fraction pseudo-replicates with higher ratio |
|---|---|---|---|---|---|
| 65 | 78 | 178 | periodic | 2.282051 | 0.434 |
| 72 | 851 | 2300 | periodic | 2.702703 | 1.000 |
| 82 | 352 | 883 | periodic | 2.508523 | 0.895 |
| 92 | 331 | 652 | periodic | 1.969789 | 0.013 |
| 101 | 264 | 549 | periodic | 2.079545 | 0.069 |
| 110 | 512 | 1324 | periodic | 2.585938 | 0.995 |
| 153 | 83 | 199 | nonper | 2.397590 | 0.615 |
| 174 | 143 | 295 | nonper | 2.062937 | 0.137 |
| 228 | 257 | 652 | nonper | 2.536965 | 0.913 |
| 389 | 373 | 767 | nonper | 2.056300 | 0.040 |
